# Muscle cell type diversification facilitated by extensive gene duplications

**DOI:** 10.1101/2020.07.19.210658

**Authors:** Alison G. Cole, Sabrina Kaul, Stefan M. Jahnel, Julia Steger, Bob Zimmerman, Robert Reischl, Gemma Sian Richards, Fabian Rentzsch, Patrick Steinmetz, Ulrich Technau

**Author notes:** These authors contributed equally to this work (Alison G. Cole, Sabrina Kaul, Stefan M. Jahnel).

## Abstract

The evolutionary mechanisms underlying the emergence of new cell types are still unclear. Here, we address the origin and diversification of muscle cells in the diploblastic sea anemone *Nematostella vectensis*. We discern two fast and two slow-contracting muscle cell populations in *Nematostella* differing by extensive sets of paralogous genes. The regulatory gene set of the slow cnidarian muscles and the bilaterian cardiac muscle are remarkably similar. By contrast, the two fast muscles differ substantially from each other, while driving the same set of paralogous structural protein genes. Our data suggest that extensive gene duplications and co-option of individual effector modules may have played an important role in cell type diversification during metazoan evolution.

**One Sentence Summary:** The study of the simple sea anemone suggests a molecular mechanism for cell type evolution and morphological complexity.

## Main Text

Animals typically consist of hundreds of different cell types, yet their evolutionary origins and relationships across organisms are still poorly understood. Moreover, most major cell type families, e.g. neurons, gland cells, epithelial cells, have often diversified into numerous related cell types, increasing the morphological complexity of the organism. Yet, the mechanisms leading to the diversification of cell types within a given species, and interspecies cell-type relationships are currently unclear and debated (*1*). Among the various cell types, muscles are one of the hallmarks of animals, which arguably played an important role in the diversification of body plans, behaviour and physiology. In the triploblastic Bilateria, which comprise the vast majority of animal species, muscles are a major derivative of mesoderm. In vertebrates for instance, three major distinct muscle types are present: striated skeletal, striated cardiac, and smooth muscles (**Fig. 1A: top**). A recent model for reconstructing cell type relationships hypothesized that related cell types will use the same core regulatory complex (CoRC; (*1*)), a collection of physically interacting transcription factors that together specify the terminal phenotype of a cell. Interestingly, despite their ultrastructural similarity, the two vertebrate striated muscles, skeletal and heart muscle, employ quite distinct regulatory complexes, whereas smooth and cardiac muscles share several of the transcription factors (*2–4*). A comparison of regulatory complexes establishing contractile cell type identity across bilaterians has hypothesized that ancestral cell types were already present in the last common ancestor of Bilateria. Two ancestral sister cell types gave rise to 1) fast contracting somatic striated myocytes and 2) slow contracting visceral smooth myocytes that further diversified into secondarily striated cardiac myocytes (*5*). Cnidarians (sea anemones, jellyfish and corals), the sister group of bilaterians, also possess contractile cells of unknown relationship to their bilaterian counterparts. Thus, to reconstruct the evolutionary relationships and diversification of muscle cell types, we investigated the transcriptional profile, anatomy and contraction abilities of muscle cells in the sea anemone *Nematostella vectensis* (Cnidaria) (**Fig. 1A: bottom, B**).

**Fig. 1.**
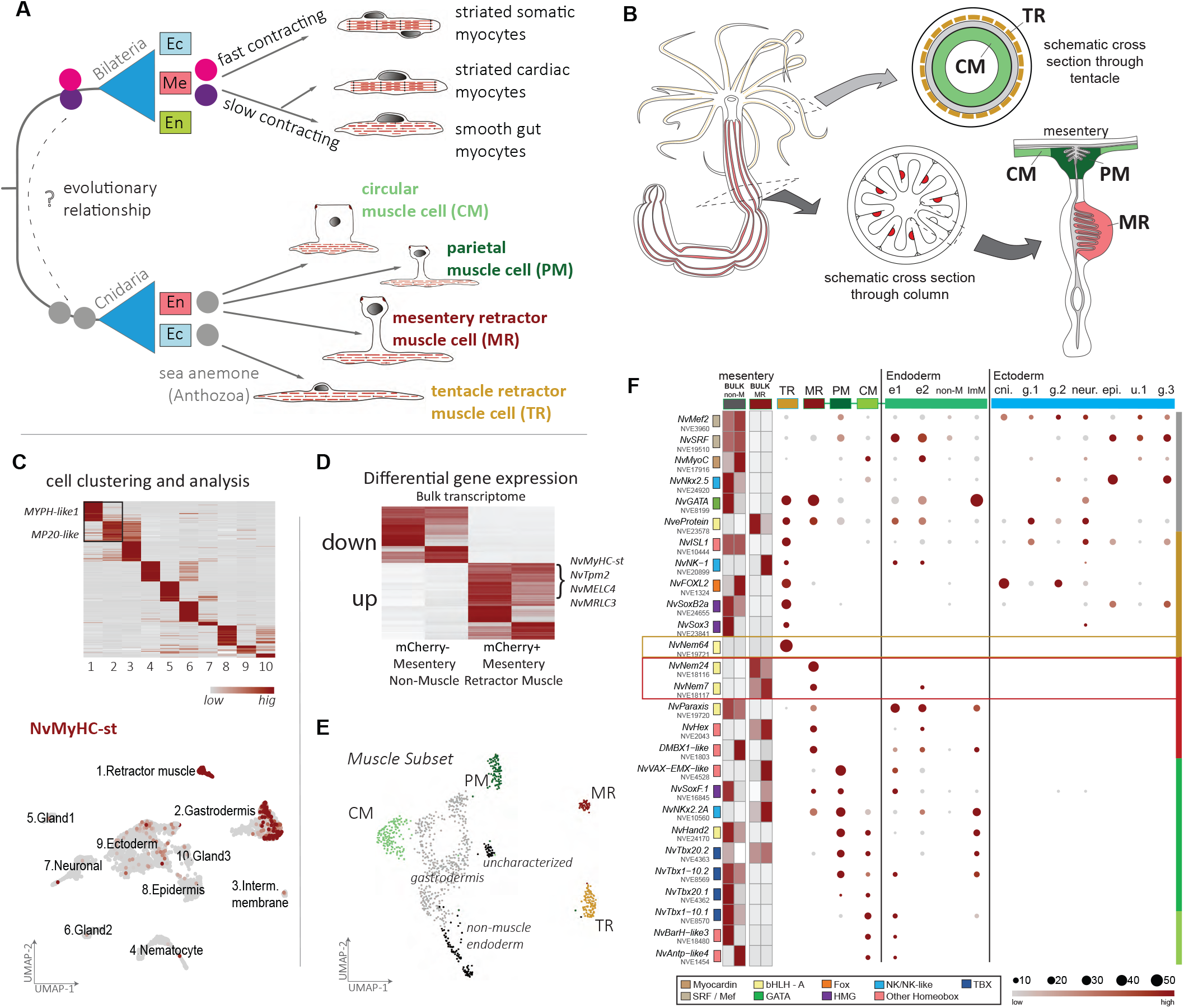
Distinct muscle cell types in the sea anemone *Nematostella vectensis*. **A** Muscle cell relationships in vertebrates (Bilateria) and the sea anemone (Cnidaria). While three major cell muscle types arise from the mesoderm in vertebrates, muscles can arise from both cell layers in the diploblastic sea anemone. Both the intra- and interspecies evolutionary relationships of these muscles are largely unclear. **B** Schematic view of *Nematostella* muscle systems and tissue used for bulk or single cell transcriptomes. The structure of the mesentery anatomy is illustrated schematically in cross section with the positions of the endodermally-derived muscles indicated: light green (CM): circular muscle; dark green (PM) parietal muscle; red (MR): mesentery retractor. **C** Heat map of differentially expressed genes across 10 identified cell populations (top). Paralogs of calponin-family genes (MYPH-like1 and MP20-like) are detected in the retractor (1) and endodermal (2) gene sets. Expression profile of *NvMyHC-st* shown on a dimensional reduction cell plot (UMAP) of the full tissue dataset (bottom). Each dot represents the transcriptome of a single cell, with expression of the gene indicated in red. Expression is detected in all cells of the retractor muscle cluster and in half of the endodermal cell cluster. Cell clusters carried forward as a data subset are indicated in the box. **D** Heatmap of differentially expressed genes of the mesentery-derived bulk transcriptomes. Average gene expression from differentially expressed genes with >2-fold change of expression between library source (mCherry negative non-muscle, 2 replicates, vs. mCherry+ mesentery retractor muscle cells, 2 replicates) are imaged. Representative muscle-related genes are up-regulated in the muscle-cell libraries. **E** Dimensional reduction cell plot (UMAP) of the muscle subset, annotated according to cluster identity. Four differentiated muscle cell clusters are identifiable, color-coded as in part (A). **F** Dot plots showing relative expression profiles of select regulatory genes. Genes in the top half of the figure illustrate the expression profile of candidate bilaterian muscle-related genes (grey bar) showing little or no expression in any of the four muscle cell types. Gene family relationships are indicated according to the colour scheme shown in the legend. Each muscle cell type is characterized by a unique combination of regulatory molecules (coloured bars). The three paralogous bHLH-family genes (*NvNem64* (ochre), and *NvNem24/NvNem7* (red) are the only regulatory factors found to be highly specific to TR and MR respectively.

Cnidaria typically have epitheliomuscular cells, i.e. with a basal contractile cellular part and an apical epithelial connection (*6*). Cnidarians are diploblastic, i.e. they develop from only two germ layers, termed ectoderm and endoderm, and epitheliomuscular cells can form from both layers. The model cnidarian *Nematostella vectensis* is organized as a solitary polyp with tentacles surrounding its mouth, opening into a blind-ending body column (*7*). The gastric cavity is compartmentalized by eight separate in-folding’s of the inner layers, called mesenteries (derived from both germ layers (*8*)). Phalloidin staining and ultrastructural studies have described five anatomically identifiable muscle sets: three longitudinally oriented muscle sets including the parietal muscles embedded within the body wall (PM), retractor muscles within the mesenteries (MR), the sub-epithelial ectodermal tentacle retractor muscle (TR), and circumferentially-oriented circular muscles of the body column and tentacles, treated here together as the endodermal circular muscle group (CM) (*9*) (**Fig. 1A,B**).

### Transcriptomic profiling reveals four molecularly distinct muscle cell populations

To characterize the molecular profile of *Nematostella* muscle cells, we used a retractor-muscle specific transgenic line that expresses mCherry under the control of a *Myosin Heavy Chain-striated type (MyHC-st)* promoter (*10*), and generated transcriptomic profiles from muscle versus non-muscle cells of the mesentery upon dissociation and Fluorescence activated cell sorting (hereon called BULK dataset). Confirming previous work (*11*), we found numerous structural muscle-related genes (e.g. *MyHC-st, myosin essential light chain (MELC)s, myosin regulatory light chain (MRLC)s, tropomyosins, calmodulins,* and *calponins*) upregulated in the library of transgenic sorted cells (**Fig. 1D**, **Extended Data 1.1**). To test whether these data are representative of all muscle types, we generated single cell (sc)RNAseq libraries from dissected juvenile tissues (**Fig. S1.1**). Analysis of the entire scRNAseq dataset revealed two cell populations that express *NvMyHC-st*, as well as many other muscle-related genes, corroborating a recently reported single cell atlas using a different scRNAseq platform (*12*) (**Fig. 1C, Extended Data 1.2**). We identified one population as retractor muscle according to overall similarity with the bulk transcriptome data (**Fig. 1C**, cluster 1; **S1.2G**). The other population was composed of cells harbouring a gastrodermal signature, including the marker *NvSnailA* (**Fig. 1C**, cluster 2; (*13*)). Analysis of this scRNAseq subset (retractor muscle + gastrodermal cells) revealed molecular profiles of 4 distinct muscle cell populations (**Fig. 1E**), forming stable clusters over a wide range of parameters (**Fig. S1.3A**). Evaluation of the spatial origin of each cell within the cluster (**Fig. S1.3B, C**) and the gene sets specific to the individual clusters (**Fig. S1.3D**) revealed correspondence with anatomically recognizable muscles: tentacle retractor muscle, mesentery retractor muscle, parietal muscle and circular muscle. The circular muscle population is distinguished from the parietal muscle by the expression of numerous collagens (marked with * in **Fig. 2A, Extended Data 1.3**), suggesting additional functions.

**Fig. 2.**
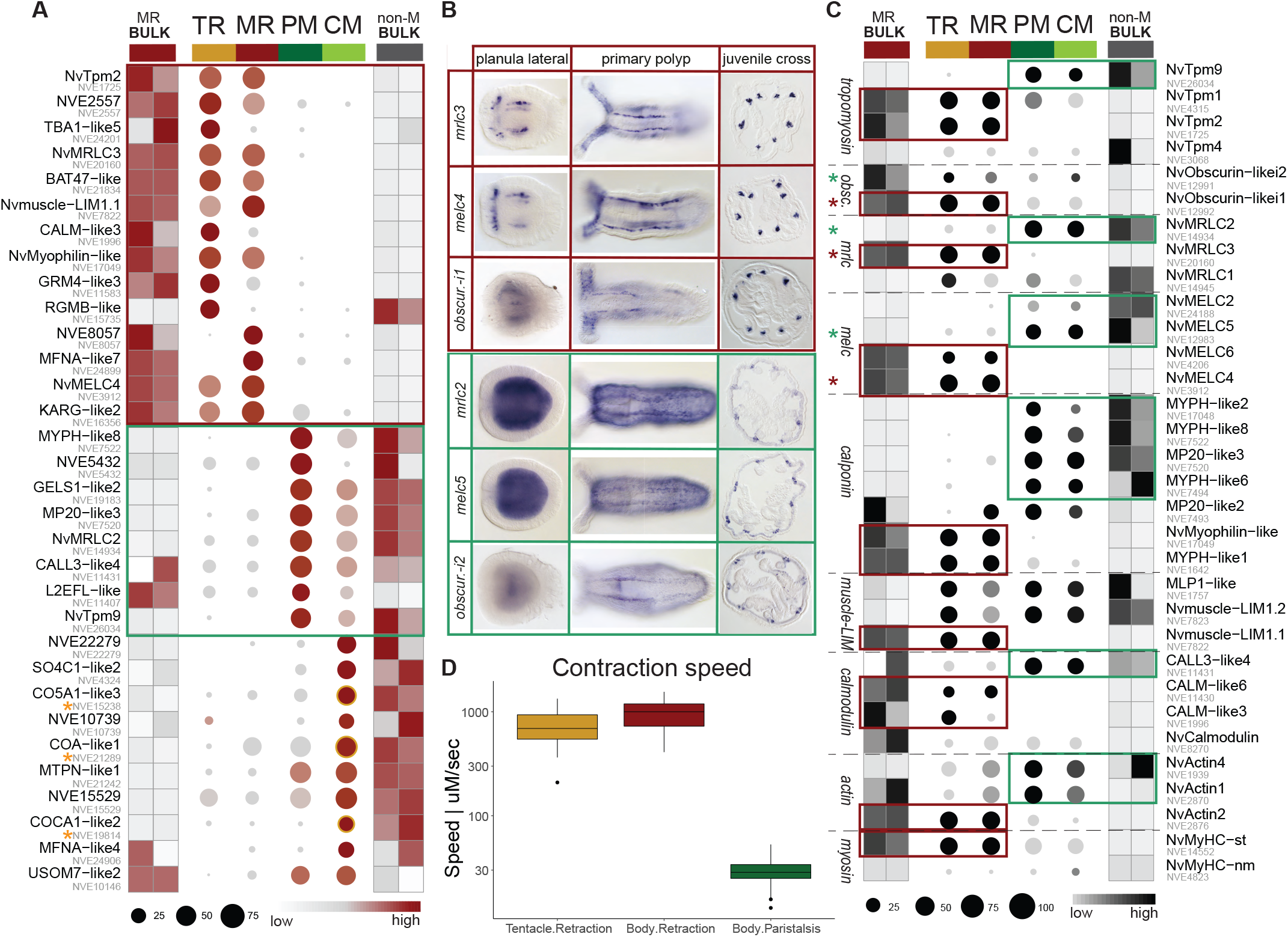
Structural protein gene expression across the four muscle cells. **A** Differential gene expression analysis identifies two distinct effector gene modules in the retractor (red) and bodywall (green) musculature, congruent with the bulk mesentery dataset. Dotplot showing expression profiles for 10 ten most significant DE genes for each muscle cluster. The size of the dot reflects the proportions of cells within the cluster that express the gene, and the colour indicates the relative expression level of expressing cells within that cluster. Cluster identity is indicated by the same colouration shown in (1E). The relative expression profile of the same gene set within the bulk dataset of the non-muscle cells (non-M BULK), and the muscle cells (MR BULK) are illustrated as square blocks. **B** in situ validation of paralogous muscle gene sets **C** Expression profile of paralogous genes illustrating differential use across clusters, set-up as in (A). Fast (red boxes) and Slow (green boxes) paralogs are highlighted. **D** Measured contraction speeds for tentacle and body column retraction (yellow and red) versus peristaltic bodywall contractions (green).

To assess the molecular basis of these four distinct cell populations, we investigated the regulatory signature associated with each cell cluster using both candidate gene (reviewed in (*14*); (*15*)) and unbiased approaches (see methods and **Fig. S1.4**). While some candidate genes commonly associated with muscle formation in Bilateria (e.g. *srf, mef2,* and *myocardin*) are detected in all four muscle cells types, they are also detected within non-muscle ectodermal derivatives at equivalent or even greater (e.g. *NvMef2*) levels (**Fig. 1F: upper grey bar**). Moreover, previous functional studies of splice variants of *NvMef2* have found no evidence for a role in the regulation of muscle development or muscle-specific genes, e.g. *MyHC-st* (*16*). Hence these genes have a much broader tissue distribution in *Nematostella,* but may have a role in muscle development in conjunction with more specific factors. Each of the four muscle populations is characterized by the expression of at least one or more highly specific regulatory molecules (**Fig. 1F, Extended Data 1.4, 1.5**), suggesting that the sea anemone *Nematostella* has diversified its muscle cell complement into 4 distinct cell types that use different regulatory complexes.

### N. vectensis *bodywall muscle regulatory signature is reminiscent of the bilaterian cardiac signature*

The bodywall (parietal and circular) muscles are most similar in their regulatory signature, commonly expressing *NvHand2, NvNkx2.2A,* two paralogs of the Tbx20 family (*NvTbx20.1, NvTbx20.2*), but differing in their use of Tbx1/10 family genes (circular: *NvTbx1/10.1*; parietal *NvTbx1/10.2*) and NK-related family genes ((*17*) *NvBarH-like3* and *NvAntp-like4* only in the CM) (**Fig. 1F**). We note that this combination of genes, together with the more globally expressed *NvSRF/MEF2* is remarkably similar to the cardiomyocyte defining set of transcription factors in bilaterians (GATA/Pannier/Serpent, Hand, NKx2.5/Tinman, Tbx5, Tbx1/10, Tbx20) (*18*). However, there are also differences: *NvTbx4/5* (derived from the common ancestor of vertebrate *Tbx4* and *Tbx5*) is expressed in parietal muscle as well as the mesentery and tentacle retractor muscles, but also broadly in the non-muscle ectoderm, thus, has a less specific expression pattern. Further, while *NvGATA* reads are detected in the parietal muscle, this gene is more strongly detected within the retractor muscles, as well as in non-muscle mesentery tissue and neuronal cells (**Fig. 1F**) ((*19*); (*8*)). *NvNk2.5* is detected in the circular muscle, but it is predominantly expressed in neural derivatives as has been previously demonstrated (*12*). However, we find an Nk2.2 ortholog (*NvNkx2.*2A), a member of the NK2/4 family, which also includes Nkx2.5, specifically expressed in all endodermal muscles (**Fig. S1.5B**). Altogether these data support homology of the bodywall muscles in *Nematostella* and the bilaterian visceral/cardiac myocyte (*8*).

### N. vectensis *retractor muscle regulatory signatures suggest convergent evolution of fast-contracting myocytes*

We next wished to investigate whether the two retractor muscle populations (MR and TR) share a set of transcription factors with the bilaterian fast contracting somatic striated myocytes. We find that these two muscle cell populations are very distinct with respect to their developmental regulators (**Fig. 1F**): the mesentery retractor muscles express *NvParaxis*, *NvDmbx*, *NvHex*, *NvNkx2.2A* and the two paralogous bHLH genes *NvNem24*, *NvNem7.* Except for *NvNem24,* these genes are also detected in other endodermal cell populations, possibly indicating the presence of muscle precursors. By contrast, the tentacle retractor muscle expresses *NvNem64*, *NvNk1*, *NvSox3,* with *NvNem64* exclusively expressed in the ectodermal muscle. Only *GATA* and *e-protein* are shared between tentacle and mesentery retractor muscles. Surprisingly, the tentacle retractor muscle in addition expresses the neuro-glandular transcription factor genes *NvSoxB2a (*aka *NvSoxB(2)), NvIsl* and *NvFoxL2*. Thus, on the level of the transcription factors the tentacle retractor cells are more reminiscent of neurons than of muscle. This is supported by the expression of ELAV, a marker of a large subpopulation of neurons in *Nematostella* (*12*) (*20*). This correlates with their anatomy: unlike the epitheliomuscular nature of the endodermal muscles, the ectodermal tentacle retractor muscles form by apical detachment from the epithelium resulting in basi-epithelial positioning, similar to ganglion neurons (*9*). Of note, *NvNem7*, -*nem24* and -*nem64* form a cnidarian-specific paralogous group with no clear orthology relationship to any of the bilaterian bHLH families (**Fig. S1.5A**) (*21*). This suggests that muscle differentiation in the sea anemone is driven by a bHLH class II:class I regulatory complex by using a mixture of conserved (*hand, paraxis*) and independently evolved factors (*NvNem7,24,64*). These bHLH transcription factors have evolved from extensive gene duplications followed by cis-regulatory sub-functionalization to define distinct classes of muscle cell types.

### N. vectensis *muscle express two classes of effector modules*

Notably, despite showing very little overlap in regulatory signature, the ectodermal tentacle retractor muscle (TR) and the endodermal mesentery retractor muscle (MR) largely share the same signature of effector molecules (**Fig. 2A, S1.2G, S2.1, Extended Data 1.6**). Similarly, we find a separate, distinct signature, shared between the parietal (PM) and circular muscle (CM) (**Fig. 2A, S1.3D, S2.1**). This is remarkable, given the ultrastructural similarity between the longitudinally oriented mesentery retractor and parietal muscles (*9*). For instance, mesentery and tentacle retractor muscle cells share expression of *NvTropomyosin2 (NvTpm2), NvObscurin-like1i1, NvMyosin regulatory light chain3 (NvMrlc3), NvMyosin essential light chain4&6 (NvMelc4&6),* the *Calponin* orthologs *NvMyophilin-like& NvMYPH-like1, NvCalmodulin-like3&6 (NvCALM-like3&6), NvMuscle-LIM1.1* and predominant usage of *NvActin2.* The bodywall musculature (circular + parietal muscle) expresses a separate set of muscle-related genes, including *NvTpm9, NvObscurin-like1i2, NvMrlc2, NvMelc2&5, NvMYPH-like2,8,6 & NvMP20-like3 (Calponin orthologs), NvCALM-like4,* and predominant usage of *NvActin1&4.* Differential expression of these variant gene sets was validated by in situ hybridization (**Fig. 2B, S2.2**). Importantly, phylogenetic analyses of these structural protein variants revealed extensive gene duplications within the cnidarian lineage in all cases (**Fig. S2.4**), independent of gene duplications occurring within the Bilateria. Plotting the structural protein genes according to gene families revealed that numerous paralogous genes are found that are specifically expressed in either TR/MR or PM/CM (**Fig. 2C**). This suggests that the four myocyte populations in *Nematostella* comprise two discernable and physiologically distinct muscle cell classes that are characterized by specific combinatorial sets of paralogous, sub-functionalized structural protein coding genes. We next asked whether the expression of these distinct paralogs of structural proteins correlates with any physiological properties of the muscle classes. Indeed, we found that the contraction speed of TR/MR class is about 50x higher than that of CM muscles (**Fig. 2D, S2.3, Extended Data 2.1,2.2**). We therefore refer to TR/MR and CM/PM as fast and slow muscle, respectively, and we presume that these variants of structural proteins convey specific properties, e.g. contractile force and speed.

### *Distinct neuronal regulation of fast contracting myocytes in* N. vectensis

Fast-contracting somatic muscles in bilaterians are stimulated at a neuromuscular junction (NMJ) by release of neurotransmitter from the presynaptic neuron that binds to appropriate receptors at the muscle cell surface. To confirm the presence of postsynaptic densities (PSD) indicative of neuromuscular junctions, we examined the retractor muscles using transmission electron microscopy (TEM) (**Fig. 3A-G**). In the tentacles, neuromuscular synapses show small clear vesicles in the terminal region of pre-synapses that are already docked to the membrane facing the synaptic cleft (**Fig. 3B, C**). The postsynaptic membrane belongs to a tentacle retractor cell and has bundles of longitudinally oriented myofilaments at the basal side of the cell. We also found synapses in the folds of the mesentery retractor cells, including terminal axons with small clear vesicles (**Fig. 3F, G**) and a PSD at the muscular post-synapse (**Fig. 3E, G**), similar to previously reported putative synapses (*22*). We were unable to find ultrastructural evidence for NMJs in the parietal and circular muscle, however, we cannot rule out that they were missed because of their sparse distribution.

**Fig. 3.**
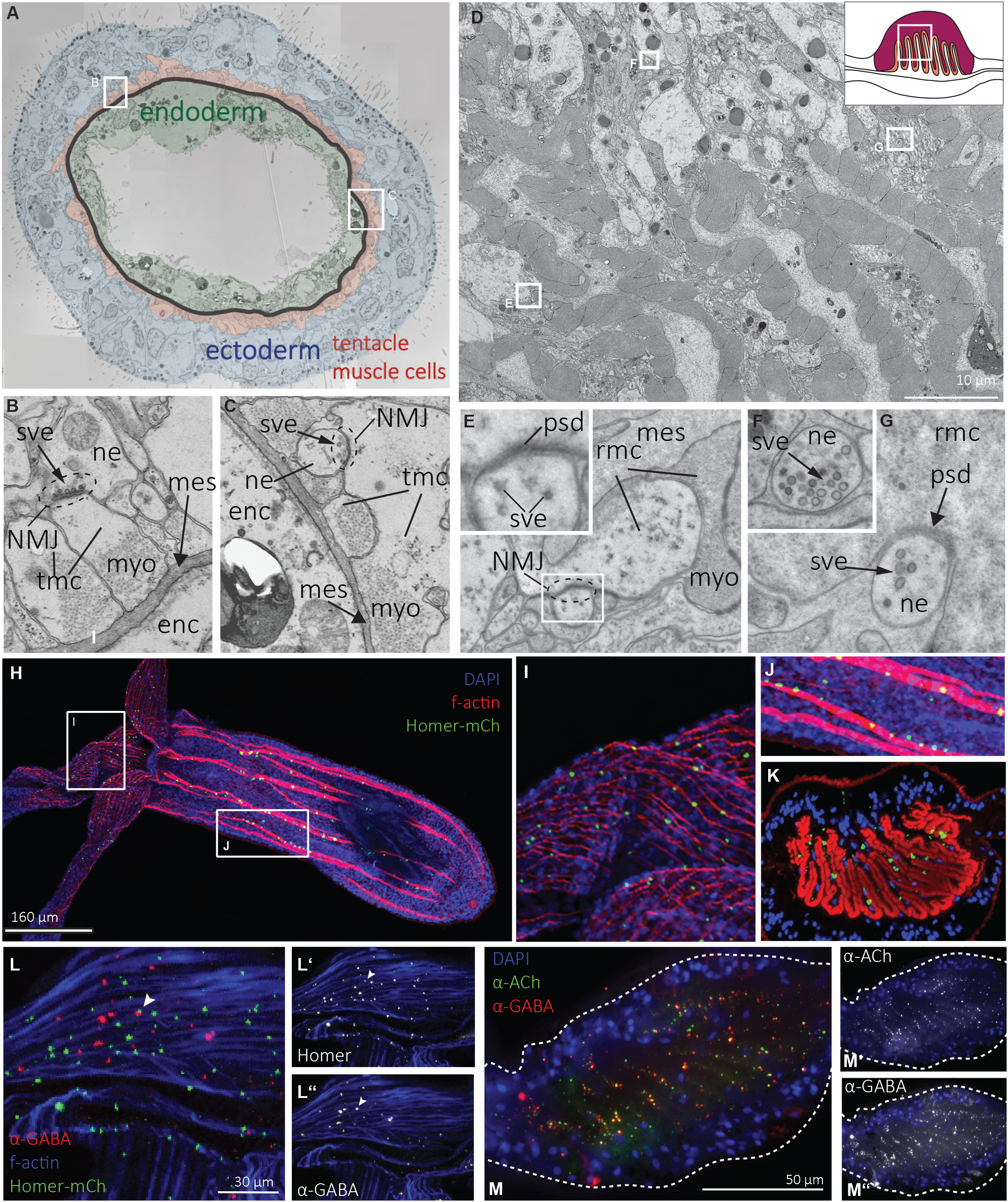
Putative neuromuscular junction and neuromuscular regulation of retractor muscles. **A** Electron micrograph of a tentacle cross section showing location of (B,C). **B, C** In several places of the tentacle retractor muscle putative neuromuscular junctions (NMJ) featuring clear vesicles (sve) are present. **D** Electron micrograph overview of the mesentery retractor approximately as indicated in the schematic **E-G** close ups marked in (**D**) showing axonal terminal ending with clear vesicles and post synaptic density (PSD) on the muscle cell. Enc: endodermal cell; tmc: tentacle muscle cell; ne: neural element; mes: mesoglea; myo: myofiliments; sve: synaptic vesicle; psd: post-synaptic density; rmc: retractor muscle cell of the mesentery. **H-M** Spot-like expression of the transgenic fusion protein Homer-mCherry (green) in the mesentery and tentacle retractor muscles suggesting putative NMJs. **H-J** Confocal images of a transgenic MyHC::Homer-mCherry primary polyp counterstained with Phallidin for F-actin. **K** Confocal image (cross section) of the adult mesentery retractor showing Homer-mCherry spots close to the muscle fibers (red), while nuclei (blue) are located at the distal end of the elongated cells. **L, M** Immunocytochemistry against GABA and Acetylcholine. **L** GABA (red) is detected in small vesicles in the tentacle ectoderm muscle (blue). Only few are located in close vicinity to Homer-mCherry (green; arrowhead). **M** Acetylcholine and GABA are detected and frequently co-localized in the folds between the mesentery retractor cells.

Ionotropic receptors are stabilized and anchored at the postsynaptic region by numerous proteins including the discs-large-, SHANK-, and Homer families (*23*). We detected transcripts for PSD proteins of the discs-large- and Homer-families within the muscle cell transcriptomes (**Fig. S3**). We reasoned that these post-synaptic proteins might also indicate NMJs in *Nematostella*. Therefore, we generated a retractor muscle-specific transgenic reporter line (*MyHC-st::homer::mCherry*) that visualizes the location of neuromuscular synapses within the mesentery and tentacle retractor muscles via a mCherry-tagged Homer protein. Spot-like structures could be detected along the tentacle retractor and mesentery retractor muscles in primary polyps, presumably indicating NMJs (**Fig. 3H-L**). In adult polyps, Homer::mCherry is localized in the elaborate folds of the mesentery retractor cells (**Fig. 3K**) in the area, where synapses were detected by TEM (**Fig. 3D-G**).

Excitatory cholinergic motor neurons that release acetylcholine into the NMJ have been found in many bilaterians including vertebrates, mollusks, annelids and echinoderms, while GABA mostly acts as an inhibitory neurotransmitter. Immuno-labelling of transgenic homer animals shows that GABA is present in small vesicles along the tentacle muscle cells, yet, they only occasionally co-localize with Homer::mCherry (**Fig. 3L-L’’**, arrowhead), indicating that this transmitter does not accumulate at the Homer-defined NMJ, but may rather reside in vesicles of neurites. GABA and Acetylcholine are detected between the folds of the mesentery retractor muscle cells, frequently in overlapping locations (**Fig. 3M-M’’**). Specific subunits of nicotinic acetylcholine receptors are detected at low levels in the scRNAseq dataset, as well as in the bulk RNAseq data, of the fast-contracting retractor muscles, but are not detected in the single cell transcriptomes of the slow-contracting bodywall muscle (**Fig. S3**). Supporting these findings, recent studies have reported the expression of several nAChRs in *Nematostella* polyps (*12*) and response of muscle contractions upon exposure of Acetylcholine (*24*). Amongst the bilaterians, only the Ecdysozoa, e.g. *Drosophila* (*25*) and crayfish (*26*) have a glutamatergic neuromuscular junction been reported. Interestingly, no transcripts of ionotropic Glutamate receptors (iGluR) could be detected in the endodermal muscles in either the BULK nor the single cell datasets (**Fig. S3**), suggesting a vertebrate-like neuromuscular regulation. By contrast, the neurotransmitter receptor repertoire in the ectodermal tentacle retractor muscle is drastically different: besides expressing multiple subunits of Acetylcholine and GABA receptors, the tentacle retractor muscle also expresses several different ionotropic glutamatergic receptor (iGluR) subunits (**Fig. S3**). While the most strongly expressed nAChR, *NvnAChRaB*, is restricted to the tentacle muscle, several of the tentacle muscle nAChRs and all iGluR are also expressed in neuronal populations (**Fig. S3**). This suggests that the neuronal regulation of tentacle retractor muscles through neurotransmitters is reminiscent of neurons and distinct from that of endodermal muscles in *Nematostella*.

### Evolutionary relationships and diversification of cnidarian muscles

Here we show that the non-bilaterian, diploblastic sea anemone has diversified its contractile cells into four muscle cell types with distinct transcriptional profiles (**Fig. 4A**). Our data provide evidence for the presence of two distinct effector modules, which correlate with ‘fast’ and ‘slow’ contractile properties. Localized neuromuscular junctions in the retractor muscles support the notion that these are fast-contracting muscles under nervous control. Sea anemone slow muscles are likely to share a common ancestry with bilaterian cardiac/smooth muscle cell type, diverged from an ancestral contractile cell expressing a core regulatory complex (CoRC) composed of members of the following transcription factor families: Nk2, Tbx1/10, Tbx20, HAND, and GATA (**Fig. 4C**). Within *Nematostella*, GATA expression is thus hypothesized to have been lost in the slow muscle, but has retained a muscle-cell expression domain within the fast-contracting musculature. By contrast, the CoRCs of *Nematostella* fast contracting muscles show little similarity with striated muscles of bilaterians, nor with the slow contracting muscle cells in *Nematostella*. Thus, in line with our previous report (*11*), we propose that fast contracting muscles, being smooth or striated, are likely to have evolved independently in cnidarians and bilaterians (**Fig. 4C**).

**Fig. 4.**
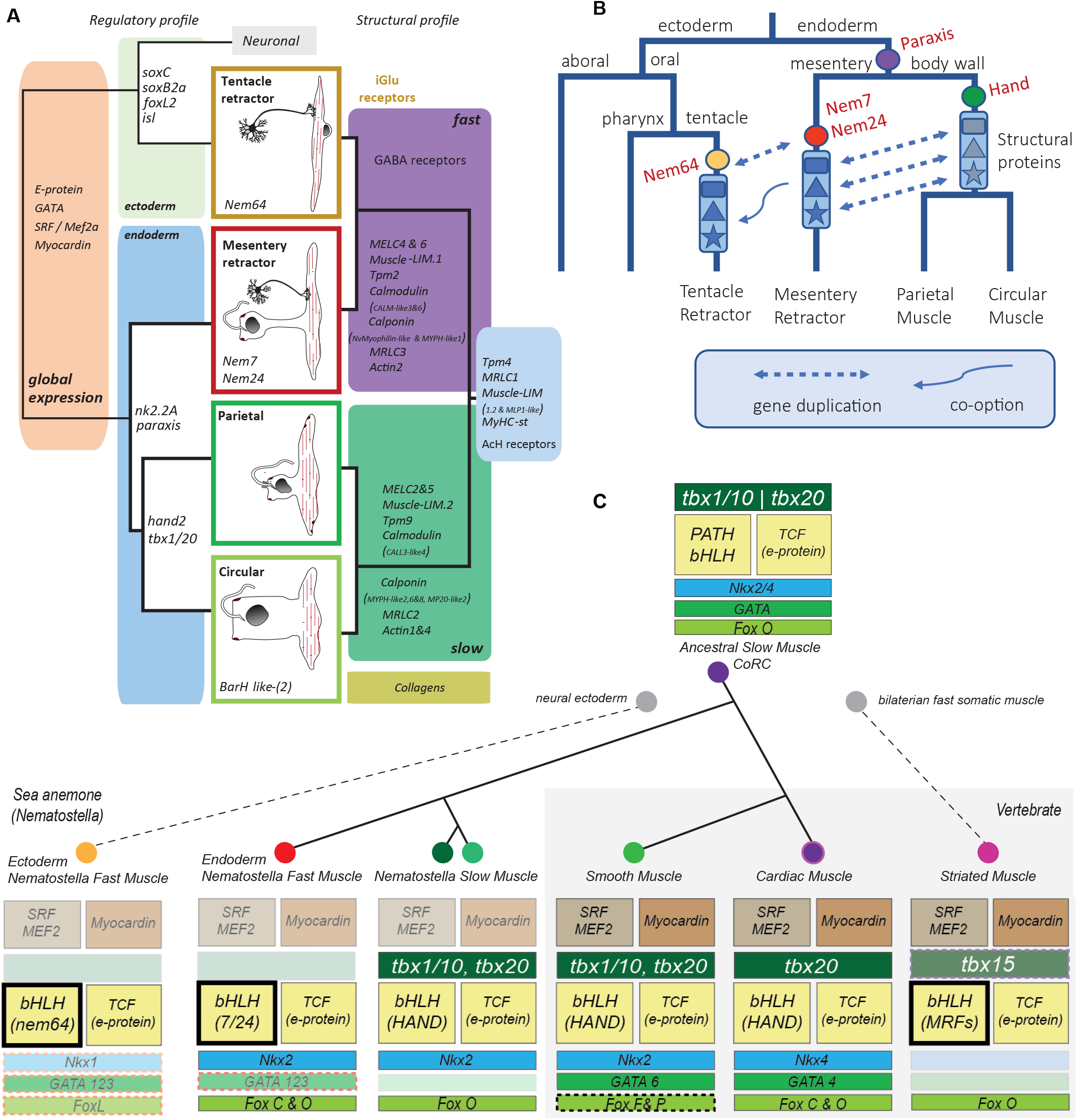
Proposed relationships and key regulatory factors underlying metazoan muscle cell evolution. **A** Non-congruence between regulatory profile and effector molecule profile among the four muscle cell types in *Nematostella*. The left side shows key regulatory genes and their association with the four muscle types shown in the center panel. The dendrogram on the left indicates the developmental relationships between the cell types. Individual genes within the cell-type boxed in the center panel are proposed here as cell-type specific regulatory factors, whereas those positioned on the dendrogram are shared between descendants of these branches. Components of the endodermal regulatory repertoire is reminiscent of the cardiac module of bilaterians (heart). On the right side, the described effector modules, or aponemes, are indicated. Note that the composition of this list is identical, but the individual paralogs are specific to each effector module. **B** Hypothesized events underlying the evolution of myocytes in the sea anemone. The fast contracting effector module is hypothesized to have evolved within the endodermal muscle populations and was then co-opted by the ectodermal epithelium after the radiation of PaTH-related bHLH proteins (**Pa** raxis, **T** wist and **H** and (see Fig S1.5)) and recruitment of one bHLH TF paralog, NvNem64, to the tentacle ectoderm within the sea anemone lineage. **C** The slow-contracting endodermal muscle in Nematostella and the slow contracting heart and smooth muscles of vertebrates share part of their regulatory set of TFs, suggesting a common evolutionary origin. By contrast, the ectodermal muscle of the tentacle, and the fast-contracting somatic musculature of vertebrates are thought to have arisen independently in the respective lineages. Regulatory molecules hypothesized to be recruited independently from related gene families are indicated in dashed boxes.

The fast and slow muscles are characterized by >25 specific paralogs coding for several effector genes, transmitter receptors and bHLH transcription factors (*Nem7, Nem24, Nem64*), strongly suggesting that the diversification of muscle cell types in this cnidarian was enabled by extensive gene duplications and sub-functionalization. In vertebrates, fast and slow twitch fibers within the somatic striated muscles are also characterized by specific expression of variants of MyHC (*27*); (*28*), suggesting gene duplications may be also instrumental in generating cellular diversity in other organisms. Interestingly, in *Nematostella*, the two fast muscles (tentacle and mesentery retractor muscle) employ distinct sets of developmental regulators, while expressing largely the same structural protein coding genes. Thus, the level of regulatory genes is not congruent with the effector genes. How could different transcription factors regulate the same battery of independent effector genes? In this context, it is striking that the two retractor muscles specifically express lineage-specific paralogs of bHLH proteins, i.e. *Nem64* in the tentacle retractor muscle and *Nem7* and *Nem24* in the mesentery retractor muscle. We postulate that *Nem64* became sub-functionalized and expressed in the outer epithelial cells of the tentacle, which also expresses several key neuronal regulators. In these cells, *Nem64* may have “hijacked” the retractor muscle specific bHLH-binding sites of the fast structural protein genes, which in the mesentery retractor muscle are bound and regulated by its paralogs *Nem7* and *Nem24* (**Fig. 4B**). bHLH transcription factors may be particularly suited for this, as they act as dimers with e-proteins, yet are rather promiscuous in their binding properties (*29*). Thus, in more general terms, paralogous transcription factors may easily co-opt even complex sets of target genes to a new cellular context facilitating diversification of cell types.

## Footnotes

1 Dataset preparation in progress. Will be released upon publication.

## Acknowledgments

We thank Michael Sixt and Alexander Leitner of the ISTA for providing us access and Doreen Milius for supervising us using the FACSAriaII machine and the Core Facility Cell Imaging and Ultrastructure Research of the University of Vienna for support. We thank Sasha Mendjan and Frank Schnorrer for critically reading the manuscript.

## Author contributions

UT conceived the study and contributed to the analyses of all data. SMJ and JS generated all bulk and sc transcriptome datasets. AGC analysed all transcriptome data with assistance from RZ. SMJ and PS performed in situ hybridization. GSR and FR generated the homer-mCherry transgenic line. SMJ and SK generated and analyzed EM and immunohistochemistry data. UT, SK, and AGC wrote the manuscript with input from all authors

## Data and materials availability

RNAseq datasets have been submitted to GEO (GSE154477), and are available^1^ for public exploration via the UCSC Cell Browser (https://cells.ucsc.edu/). All other data is available in the main text or the supplementary materials.

## Supplementary Materials

