## Supplemental material for "Muscle cell type diversification facilitated by extensive gene duplications"

**This PDF file includes:**

Materials and Methods
Figures S1-S3
References (S1-18)

**Other Supplementary Materials for this manuscript include the following:**

Movies Extended Data
2.2 External Data S1-S5

### Materials and Methods

Bulk transcriptome data processing and analysis

In order to enrich for muscle tissue in *N. vectensis* we used the muscle specific reporter line MyHC::mCherry line (*1*), which specifically expresses mCherry in the retractor and tentacle longitudinal muscle of the polyp. Adult polyps (>4cm length in a relaxed state) with strong mCherry expression in the retractor muscle region were chosen and relaxed for 30 minutes by adding a few drops of 7% $MgCl_2$ to a large petri dish. Subsequently animals were pinned down with needles, the body wall was opened from pharynx to the foot. After lateral fixation of the opened bodywall with additional pins, as much mesenterial tissue as possible was removed distal from the retractor muscle. Tissue enriched for $mCh^+$ retractor muscle was then removed by cutting through the intermuscular region and pooled resulting in two replicates with muscle tissue derived from eight animals in each case. Tissue was dissociated similarly as described previously (*2*) by adding dissociation mix (50µl papain (Sigma-P4762; 3.75mg/ml in *Nematostella* medium (=1/3 artificial sea water), 50µl Collagenase (Sigma-C9407; 1000U/ml in *Nematostella* medium), 3.5µl 0.1M DTT) to at most 4 retractor muscles at once in a 1,5ml Eppendorf tube and left at room temperature overnight (12-15h). Next morning tubes were flicked gently and successful dissociation into single cells was examined under the microscope. Afterwards cells were spun down (300G, 15min) and the pellet was washed with *Nematostella* medium and resuspended carefully. This process was repeated twice and dissociated and washed cells were combined and stored at 4°C until further processing (<1h). Cells were sorted using FACSAriaII and collected in a 15ml falcon tube containing TRIzol LS Reagent (Thermo Fisher Scientific), maintaining a final dilution of 3:1. In order to prevent degradation sorted cells were mixed with TRIzol regularly. Samples were kept on ice and total RNA was extracted according to the manual. Only samples with a RQI>7.3 were accepted for library preparation (NEB polyA) and subsequently sequenced (HighSeq 2500) single-end with a read length of 50.

Raw reads were processed using cutadapt (*3*) for adapter trimming and SortMeRna (*4*) in order to remove ribosomal RNA. Processed reads of *N. vectensis* were subsequently mapped to the repeat-masked genome using tophat (*5*) and reads with a mapping quality <20 were discarded. Aligned reads were counted with HTSeq (*6*) and differential gene expression analysis was performed using edgeR (*7*). Workflow is depicted in **Figure S1.1F**.

Single cell RNA sequencing

5-month old polyps were dissected into 4 separate tissues: tentacles, mesenteries, pharynx, bodywall (see **Figure S1.1A:E**). Tissue pieces were treated with undiluted TrypLE$^{TM}$ Select (Thermo Fisher, A1217701). For 1 hour the tissues were allowed to disintegrate in the enzyme with gentle agitation on a shaker, then dissociation into single-cell suspensions was completed with occasional pipetting over the course of another 1.5 hrs. Cell viability and counts were assayed with a Cellometer X2 (Nexcelom), and suspensions were diluted to 1000 – 1700 cells/uL (**Figure S1.1G**). Cell suspensions were kept on ice for no more than 1 hr, and loaded into a 10x Genomics single cell platform using Vs.2 reagents. Libraries were generated following the manufacturers protocol. Sequenced libraries were processed through the cellranger 2.1.0 pipeline using default parameters and forcing the pipeline to recover 7000 cells for each library. Reads were mapped to a customized transcriptome (*8*), wherein all gene models were extended by 1000bp in the 3' direction, or until the start of the next gene model in the same orientation.

Single cell transcriptomic analysis

The resulting count matrices were imported in R for further processing using the R-package Seurat Vs3 ((*9*); (*10*)). The four libraries were merged using the merge function, and the resulting dataset was then filtered for cells containing at least 100 genes, and outliers with UMI counts >30 000 were removed, as these likely reflect cell multiplettes (**Figure S1.1G**). To facilitate inter-library comparisons, gene expression values were first standardized within each library, and these relative expression values were used for dimensional reduction and cell clustering. Data was scaled to 5000 reads, log normalized, and 2000 variable genes were identified using the FindVariableGenes function (**Figure S1.2A**). Reduction algorithms were applied to the dataset (principle components analysis, UMAPs (*11*)), and hierarchical clustering was performed using the top 20 identified principle components (**Figure S1.2B**). Cluster-specific genes sets were determined for each cell population using the Seurat FindAllMarkers function using all variable genes, requiring genes to be expressed in at least 10% of the cell population and showing a log fold change in expression of at least 1 for variable genes, or 0.6 for transcription factors specifically, and a p-value of less than 0.0001. The resultant gene lists were examined in order to assign a population identity to each cluster (**Figure S1.2D**). Muscle clusters were identified based on the presence of some pan-muscle markers and a subset of the full dataset was processed in a similar manner. For the muscle-only subset, variable genes were again determined, dimensional reduction algorythms were applied, and

10 principle components were used for hierarchical clustering. Assessment of gene usage across clusters is visualized in terms of number of cells detected and average expression levels using the DotPlot function (**Figure S1.4A**). Specific structural genes were identified as having at least 20 reads within the cluster, and absent from the ectodermal portion of the dataset (<200 reads) (**Figure S2.1**). In the case of transcription factors and channel proteins, were overall detection was low, >5 reads associated with genes of interest within the cell population is taken as evidence of expression within that population as a whole. Muscle-specific genes were selected as being absent from non-muscle cell populations within the full dataset (<25 reads) (**Figure S1.4B**). The R script for generating the data objects, analysis, and figures is provided as **Extended Data S3**.

MyHC::homer-mCherry transgenic line

The MyHC::homer-mCherry line was generated by meganuclease-assisted transgenesis ((*12*) (*13*)). The open reading frame of a *Nematostella* homer homolog (NCBI XP_001637685) was cloned with AscI restriction sites into the pMyHC::mCherry vector to generate pMyHC::homer-mCherry. After digestion with I-SceI, the plasmid was injected into fertilized eggs at a concentration of 20ng/µl. Offspring of the injected animals was screened for germline transmission.

Phylogenetic analysis

Sequences used for phylogenetic analyses were either downloaded from UniProt (http://www.uniprot.org) or NCBI (https://www.ncbi.nlm.nih.gov) databases. Alignment was done via MAFFT (*14*) and maximum likelihood trees were calculated using IQ-TREE (*15*). Neighbour-joining trees were generated with ClustalX (*16*). Phylogenetic trees were subsequently visualized with FigTree v1.4.3. and processed with Adobe Illustrator. Sequence alignments are provided (**Extended Data S4**)

Calculation of muscle retraction speeds

Juvenile polyps (4-8 tentacles) of the MyHC::mCherry transgenic line (*1*) were used for all experiments. Animals were imaged with Nikon Eclipse TS100 equipped with a Nikon DS-Qi camera. The exposure and gain were set as short as possible to ensure highest possible frame rate and sharp images. The animal was placed within frame and videos were recorded before and after a dilution of Acetic-Acid 1:1000 in ELIX was used to trigger tentacle contraction whereas a ratio of 1:100 was used to trigger a full body contraction. Videos were then processed with the imaging software ImageJ. The distance between the beginning and the end of the fluorescent (mCherry) mesentery retractor were measured frame by frame, and the time point showing the greatest distance was used to calculate contraction speed of the body column as a proxy for mesentery contraction rate. Similar measurements were made from the base to the tip of the tentacles in order to estimate tentacle retractor speed. To calculate peristaltic contraction images were stabilized using the TrakEM2 plugin for Fiji; landmarks were used to measure the maximum and minimum circumference of the animals. Contraction speed was then calculated by multiplying the measured values Pi. See **Figure S2.3** for illustration.

Identification of receptor subunit orthologs

*N. vectensis* genes for ionotrophic receptor subunits have been identified by searching the publicly available genome (*17*) for specific criteria. Ionotrophic glutamate subunits were filtered with HMMTOP for having at least three transmembrane domains, a minimum length of 500 aa, and the Pfam domains PF00060 and PF01094. Ionotrophic GABA and Acetylcholine receptor sequences have been retained, when they passed filtering for Pfam02931 (Neur_chan_LBD), at least four transmembrane membrane domains and a minimum length of 400 aa. Further allocation to either GABA or acetylcholine receptors has been done by using BLAST.

Fixation, whole-mount in situ hybridization and imaging

Animals used for in situ hybridization were treated as described previously (*18*). Stained pieces of juveniles were either prepared for cryosectioning (MRLC1, MRLC 2 MRLC,3, MELC1, MELC 2, MELC 4, MELC 5, MELC 8; Obscurin-like3) as described (*18*), or treated as follows: Samples were infiltrated with 10% gelatine in PBS at 37°C for 30 min. After that samples were transferred to a mold filled up with liquid gelatine solution and subsequently orientated longitudinally. Solidified blocks were then post-fixed in 3.7% formaldehyde at 4°C overnight, washed in PBS and sectioned at 20-30µm using a Vibratome Leica VT 1200S. Sections were mounted on slides with 86% glycerol, visualized and imaged on a Nikon 80i upright microscope. Images were processed (cropping, level adjustment) using Adobe Photoshop CC15. All figures were assembled and schematics drawn in Adobe Illustrator CC15.

Phalloidin staining, immunolabelling and visualization

*Nematostella* primary polyps of the *MyHC::mCherry, MyHC::homer-mCherry,* and wildtypes were relaxed in 0.7% $MgCl_2$ in *Nematostella* medium for 30 min and subsequently fixed in 3.7% formaldehyde in PBT (PBS, 0.4% Tween) at 4°C overnight and washed thoroughly the next day. For antibody (AB) staining of the *MyHC::homer-mCherry* and non-transgenic polyps and adults, samples were processed the same day after 3 hours of fixation, blocked 2 hours in blocking solution (20% sheep serum, 1% BSA in PBT) and subsequently incubated into first AB solution containing α-Acetylcholine (MerckMillipore # AB5522, 1:200), α-GABA (Sigma #A0310, 1: 250) in PBT over the weekend. Samples were then rinsed three times in PBT and incubated in secondary ABs in PBT (AlexaFluor 488 goat anti-rabbit, Thermo #A-11008 and AlexaFluor 568 goat anti mouse, Thermo #A-11019, each 1:500) over night. Samples were then incubated in Phalloidin-AlexaFluor 488 (Thermo # A12379) (3µl/100µl PBT) and DAPI (1:1000 in PBT) over night at 4°C in the dark, washed and mounted on a glass slide in VECTASHIELD Antifade Mounting Medium and imaged on a Leica SP5 confocal microscope.

Transmission electron microscopy

All samples were anesthetized with 0.7% MgCl2 and put on ice (primary polyps, adults) and then fixed with 2.5% glutaraldehyde in 0.1 M cacodylatebuffer (pH 7.2) for 1 h (planulae, primary polyps) or overnight (adults) at 4°C. After fixation, samples were either stored in 1.25% glutaraldehyde in 0.1 M cacodylate buffer (pH 7.2) at 4°C or processed immediately. Subsequently, they were rinsed in the same buffer used for fixation. Samples were postfixed in 1% OsO4 in 0.1 M cacodylate buffer (pH 7.2) for 30 min and washed with 0.1 M cacodylate buffer (pH 7.2). Thereafter they were dehydrated through a graded series of ethanol and acetone and embedded into Low Viscosity Resin (Agar) following sectioning using standard techniques. After staining with 2% uranylacetate (20 min) and lead citrate (10 min), sections were examined with a Zeiss Libra 120 transmission electron microscope.

Figure S1.1

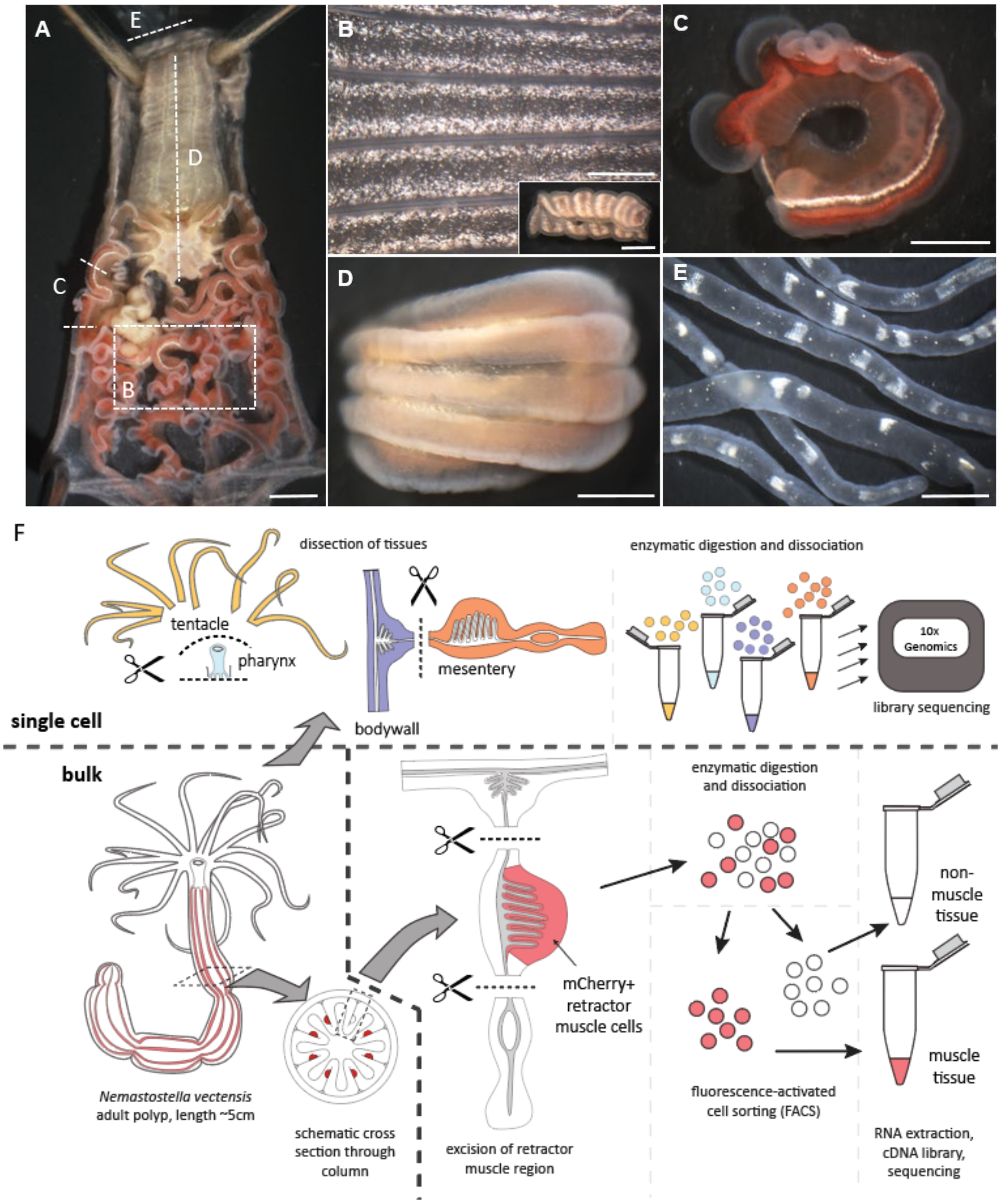


G

| | Diluted | | |
|---|---|---|---|
| | Concentration | Viability | |
| | total cell number, million cells/ml | % | Pass filter |
| **Bodywall** | 1.4 | 76.5 | 1079 |
| **Mesentery** | 1.48 | 74.9 | 583 |
| **Pharynx** | 1.03 | 85.3 | 1720 |
| **Tentacle** | 1.35 | 85.5 | 868 |

H

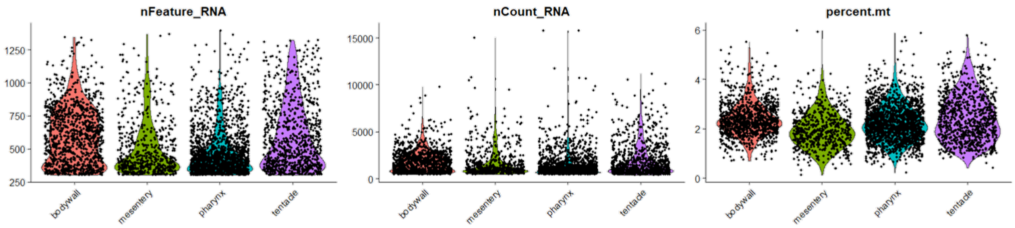

**Figure S1.1**:
**A** Dissection procedure. First, tentacles were cut off close to the tentacle base, indicated with the dashed line at the very top (E). Then all mesenteries were removed in order to expose the body wall. A small piece of a single mesentery, outlined by dashed line (C) was collected for dissociation. The pharynx was isolated and longitudinally cut in two halves. Only one half was used for library preparation (D). Finally, a rectangular piece of tissue was excised from the body wall, the inset shows its outline. (B). All scale bars 500 µm. **B** Body wall. Inset shows the outline of the tissue piece. The horizontal lines correspond to the parietal muscle. **C** Mesentery. The retractor muscle is the brown tissue in the center. **D** Pharynx. **E** Tentacle. **F** Schematic representation of transcriptome generation. Isolated tissues described above were dissociated and processed with a 10xGenomics Chromium single cell controller (top). To generate the bulk transcriptomes, the mesentery was first removed from an MyHC-st::mCherry transgenic animal, and the central retractor-muscle containing piece was dissociated. Cell suspensions were FACS sorted into fluorescent (muscle) and non-fluorescent (non-muscle) fractions. RNAseq libraries were generated from each isolated cell fraction in bulk. **G** Cell yield and viability estimates from the tissue dissociations processed as single cells, including recovered high quality single cell transcriptomes from each library (pass filter). **H** Cells from each library show similar recovered numbers of genes detected (nFeature_RNA), transcripts (nCount_RNA) and mitochondrial fraction (percent.mt).

Figure S1.2

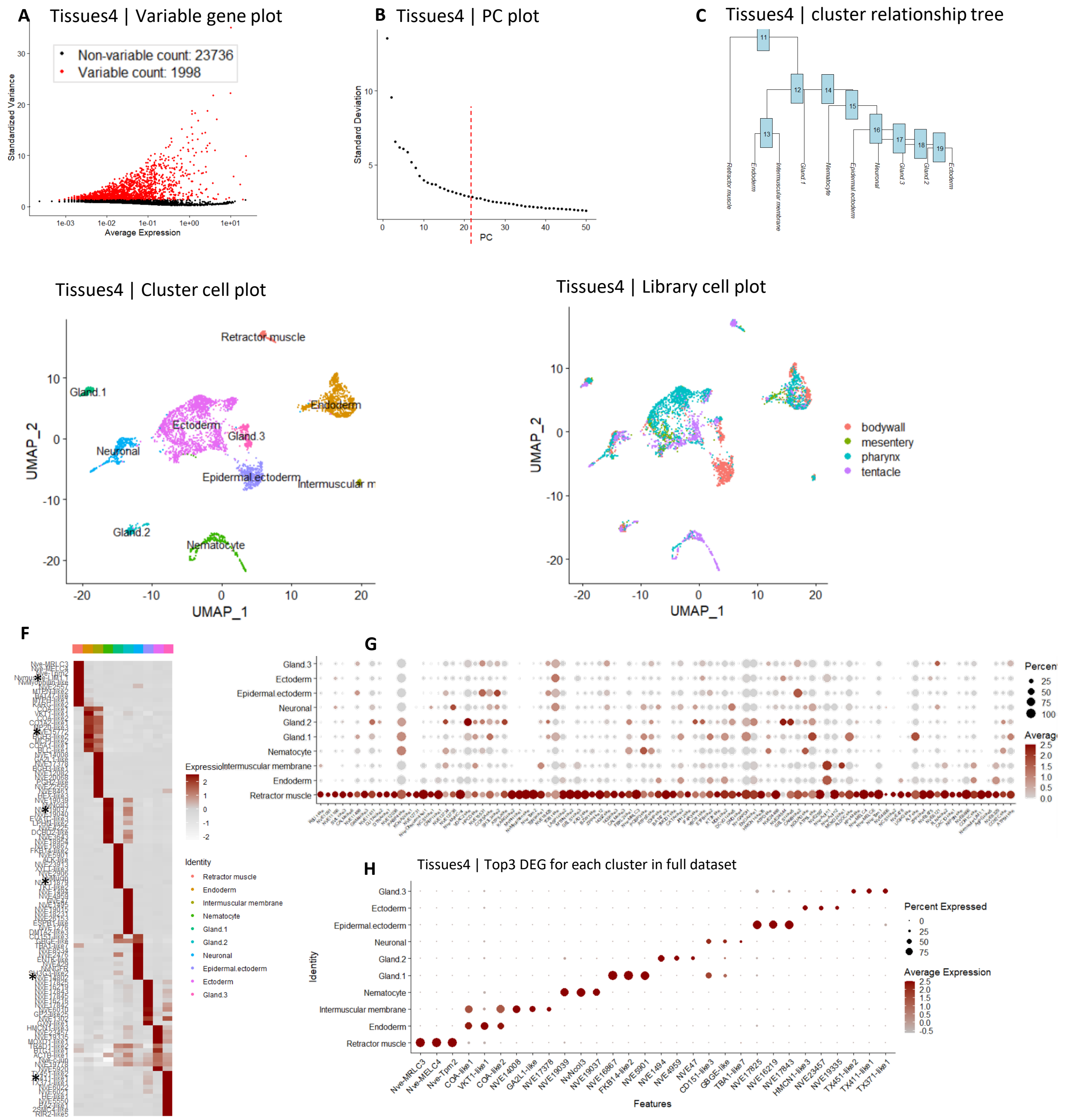
A
Tissues4 | Variable gene plot
Non-variable count: 23736
Variable count: 1998
Standardized Variance
Average Expression
B
Tissues4 | PC plot
Standard Deviation
PC
C
Tissues4 | cluster relationship tree
Retractor muscle
Endoderm
Intermuscular membrane
Gland.1
Nematocyte
Epidermal ectoderm
Neuronal
Gland.3
Gland.2
Ectoderm
Tissues4 | Cluster cell plot
Retractor muscle
Gland.1
Endoderm
Ectoderm
Gland.3
Neuronal
Epidermal.ectoderm
Intermuscular m
Gland.2
Nematocyte
UMAP_1
UMAP_2
Tissues4 | Library cell plot
bodywall
mesentery
pharynx
tentacle
F
Expression
Identity
Retractor muscle
Endoderm
Intermuscular membrane
Nematocyte
Gland.1
Gland.2
Neuronal
Epidermal.ectoderm
Ectoderm
Gland.3
G
Gland.3
Ectoderm
Epidermal.ectoderm
Neuronal
Gland.2
Gland.1
Nematocyte
Intermuscular membrane
Endoderm
Retractor muscle
H
Tissues4 | Top3 DEG for each cluster in full dataset
Identity
Features
Percent Expressed
Average Expression
Nve-MRLC3
Nve-MELC4
Nve-Tpm2
COA-like1
VKT1-like1
COA-like2
NVE14008
GA2L1-like
NVE17378
NVE19039
NvNcol3
NVE19037
NVE16867
FKB14-like2
NVE5901
NVE1494
NVE4959
NVE47
CD151-like3
GBGE-like
TBA1-like7
NVE17825
NVE16219
NVE17843
HMCN1-like3
NVE23457
NVE19335
TX451-like2
TX411-like1
TX371-like1

**Figure S1.2**;
Processing of the tissue dataset with the Seurat Vs3 package in R. **A** Variable gene plot, genes used are highlighted in red **B** Principle component plot, red line indicated the cut-off for included PCs. **C** Resultant hierarchical clustering tree. **D, E** UMAP dimensional reduction cell plot colored by labeled cell clusters (E), or library of origin (D). **F** Average expression profile of the top 10 differentially expressed genes from each cluster as in **Figure 1C (top)**. Cells identified as the intermuscular membrane show clear overlap with the endodermal cluster, however expresses a distinct repertoire of genes. Similarly, uncharacterized Gland.2 population shares expression with Nematocytes. **G** Expression profile of the set of the genes upregulated in the BULK retractor muscle dataset that are represented by at least 50 reads in the single cell dataset, plotted as a DotPlot on the single cell dataset. This gene set clearly identifies a retractor muscle cell cluster. **H** DotPlot of the expression profile of the top three differentially expressed genes for each cluster of cells.

Figure S1.3

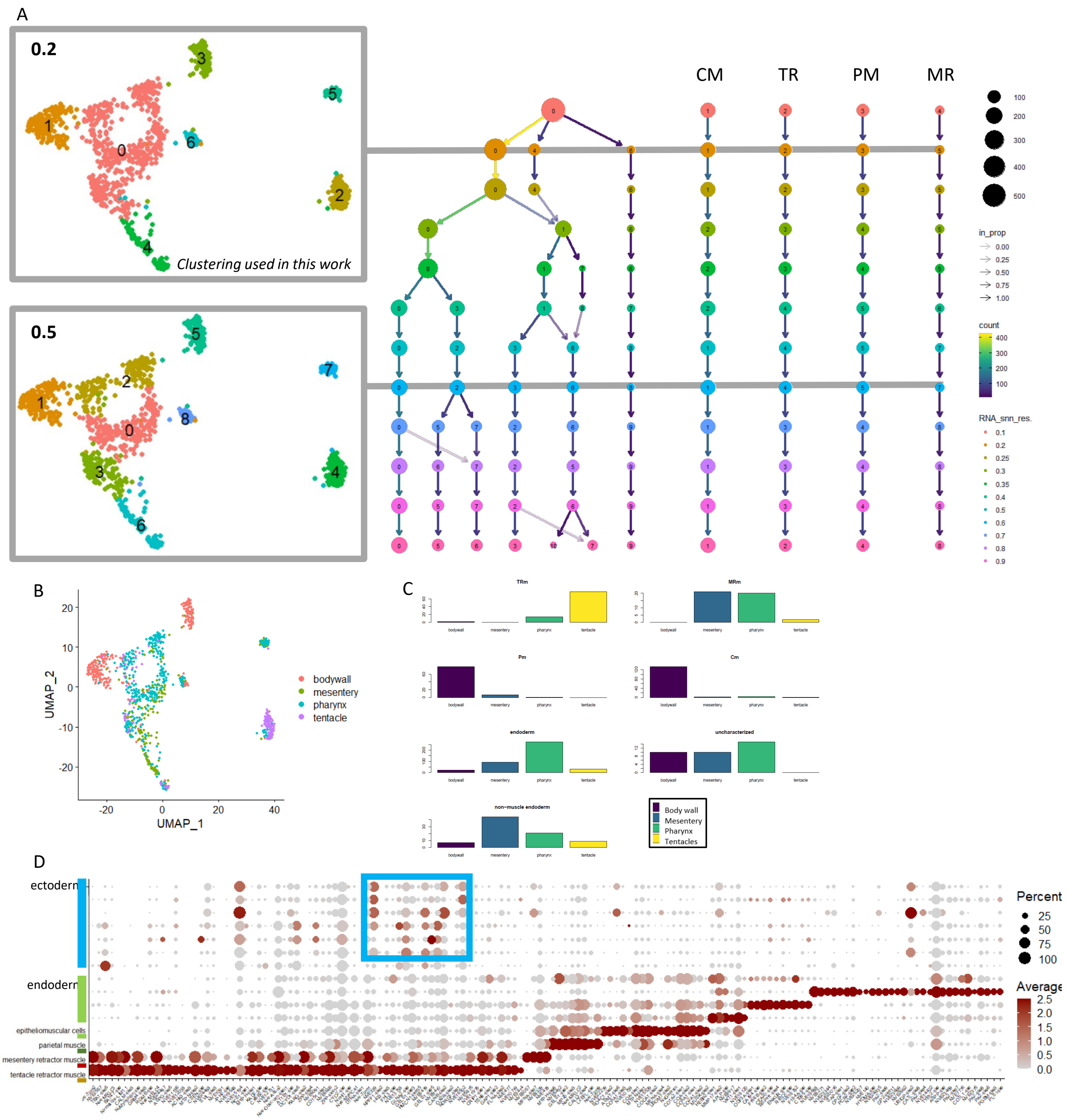
A
0.2
Clustering used in this work
0.5
CM
TR
PM
MR
in_prop
count
RNA_snn_res.
B
UMAP_2
UMAP_1
bodywall
mesentery
pharynx
tentacle
C
TRm
MRm
Pm
Cm
endoderm
uncharacterized
non-muscle endoderm
Body wall
Mesentery
Pharynx
Tentacles
D
ectoderm
endoderm
epitheliomuscular cells
parietal muscle
mesentery retractor muscle
tentacle retractor muscle
Percent
Average

**FigureS1.3**:
Processing of the muscle/endodermal subset. **A** Distribution of cells within identified clustering at increasing resolutions. The four identified muscle populations are stable at all cutoff points (resolution). **B** UMAP dimensional reduction cell plot coloured by library of origin. **C** distribution of cells from each library across the seven cell clusters used in all analyses (res = 0.2). **D** Expression profiles of all differentially expressed genes from each muscle subset cluster of cells, plotted across the entire dataset. The largest set of DEGs belongs to the ectodermally derived tentacle retractor muscle. Many of these genes are shared with either the mesentery retractor (MR), or in some cases with other ectodermal derivatives (box)

Figure S1.4 :

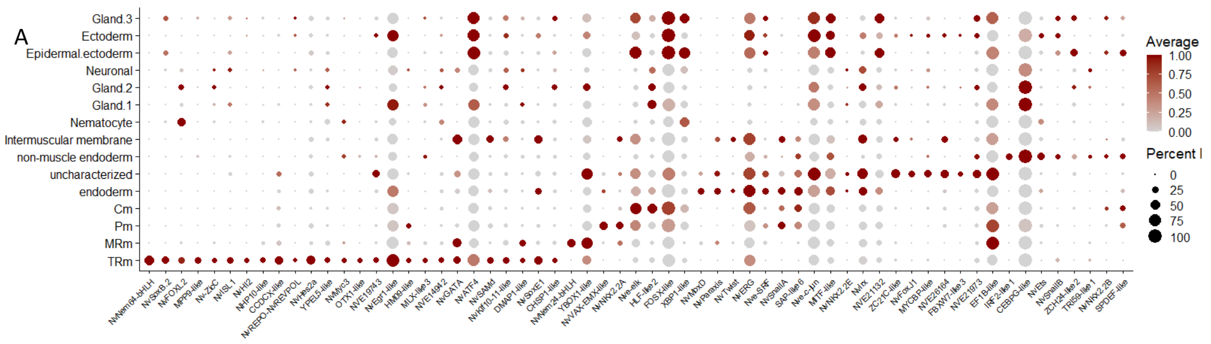


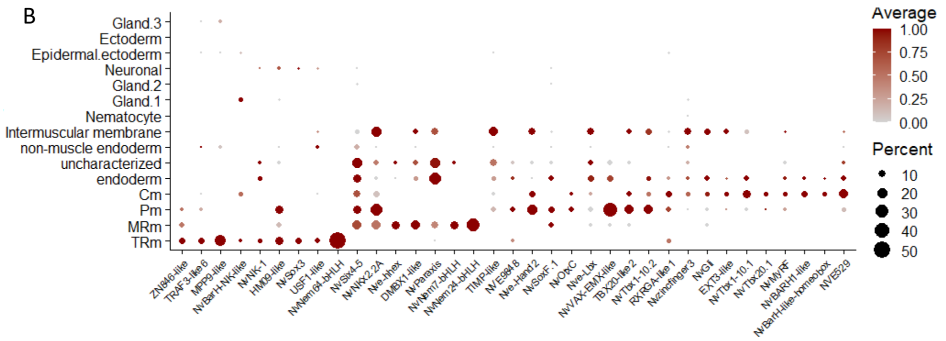


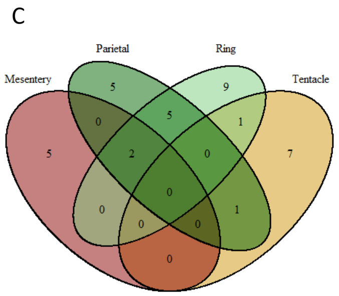


D

*SRF,'Nve-MyoC',Nk,FKH,GATA,SOX,TBX,Homeodomain with >5 reads in any muscle:*

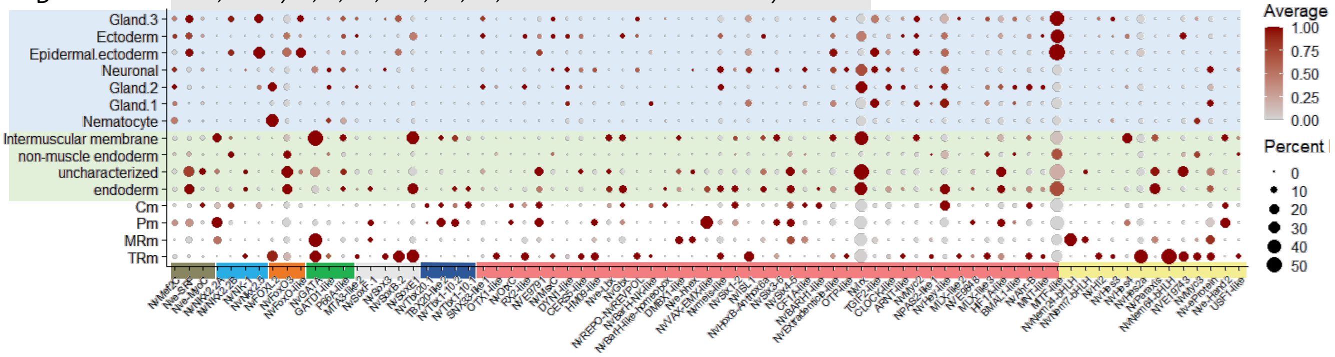


### Supplementary files: Gene lists

| **Ext.Data1.5** | **Ext.Data1.6** | **Ext.Data3** |
|---|---|---|
| ExtendedDataMUSCLE_DEG_TFs | ExtendedDataMUSCLE_AllTfsFiltered | Gene annotations |

**Figure S1.4:**

Expression profiles of transcription factors. **A** Differentially expressed transcription factors as calculated with the Seurat 'FindMarkers' function, restricting the genes set to only those with DNA-binding motifs. **B** All detectable transcription factors from each differentiated muscle cluster with fewer than 25 reads within the ectodermal cell clusters of the full dataset. **C** VennDiagram of overlapping genes of the transcription factor set. **D** all putative orthologs of transcription factor families with known roles in muscle development in bilaterians. Putative members of the gene families were filtered for expression of at least 5 reads within any muscle cell population.

Figure S1.5

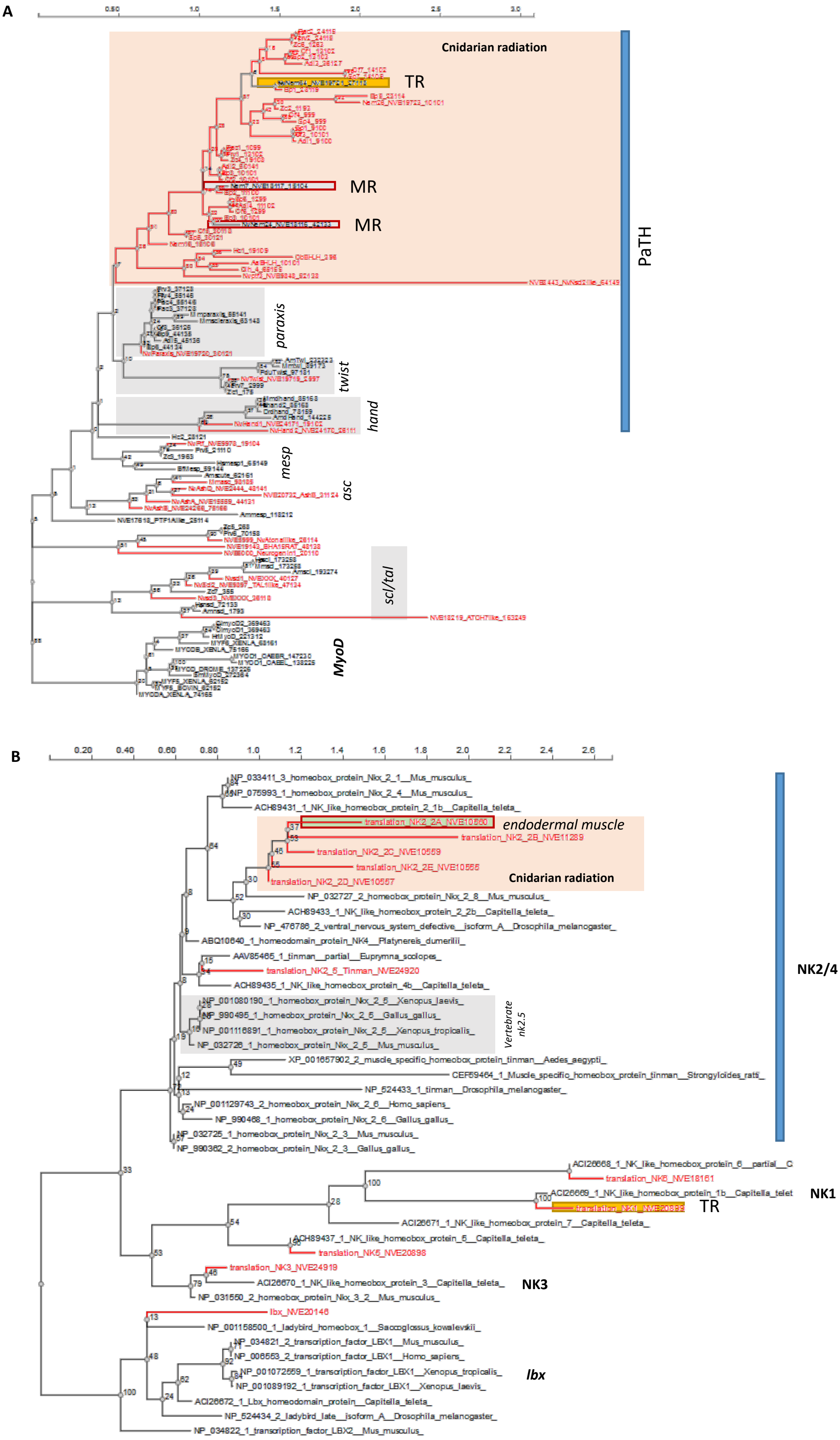

A
Cnidarian radiation
TR
MR
MR
PaTH
paraxis
twist
hand
mesp
asc
scl/tal
MyoD
B
endodermal muscle
Cnidarian radiation
NK2/4
Vertebrate nk2.5
NK1
TR
NK3
lbx
NP_033411_3_homeobox_protein_Nkx_2_1__Mus_musculus_
NP_075993_1_homeobox_protein_Nkx_2_4__Mus_musculus_
ACH89431_1_NK_like_homeobox_protein_2_1b__Capitella_teleta_
translation_NK2_2A_NVE10560
translation_NK2_2B_NVE11289
translation_NK2_2C_NVE10559
translation_NK2_2E_NVE10556
translation_NK2_2D_NVE10557
NP_032727_2_homeobox_protein_Nkx_2_8__Mus_musculus_
ACH89433_1_NK_like_homeobox_protein_2_2b__Capitella_teleta_
NP_476786_2_ventral_nervous_system_defective__isoform_A__Drosophila_melanogaster_
ABQ10640_1_homeodomain_protein_NK4__Platynereis_dumerilii_
AAV85465_1_tinman__partial__Euprymna_scolopes_
translation_NK2_5_Tinman_NVE24920
ACH89435_1_NK_like_homeobox_protein_4b__Capitella_teleta_
NP_001080190_1_homeobox_protein_Nkx_2_5__Xenopus_laevis_
NP_990495_1_homeobox_protein_Nkx_2_5__Gallus_gallus_
NP_001116891_1_homeobox_protein_Nkx_2_5__Xenopus_tropicalis_
NP_032726_1_homeobox_protein_Nkx_2_5__Mus_musculus_
XP_001657902_2_muscle_specific_homeobox_protein_tinman__Aedes_aegypti_
CEF59464_1_Muscle_specific_homeobox_protein_tinman__Strongyloides_ratti_
NP_524433_1_tinman__Drosophila_melanogaster_
NP_001129743_2_homeobox_protein_Nkx_2_6__Homo_sapiens_
NP_990468_1_homeobox_protein_Nkx_2_6__Gallus_gallus_
NP_032725_1_homeobox_protein_Nkx_2_3__Mus_musculus_
NP_990362_2_homeobox_protein_Nkx_2_3__Gallus_gallus_
ACI26668_1_NK_like_homeobox_protein_6__partial__C
translation_NK6_NVE18161
ACI26669_1_NK_like_homeobox_protein_1b__Capitella_telet
translation_NK1_NVE20893
ACI26671_1_NK_like_homeobox_protein_7__Capitella_teleta_
ACH89437_1_NK_like_homeobox_protein_5__Capitella_teleta_
translation_NK5_NVE20898
translation_NK3_NVE24919
ACI26670_1_NK_like_homeobox_protein_3__Capitella_teleta_
NP_031550_2_homeobox_protein_Nkx_3_2__Mus_musculus_
lbx_NVE20146
NP_001158500_1_ladybird_homeobox_1__Saccoglossus_kowalevskii_
NP_034821_2_transcription_factor_LBX1__Mus_musculus_
NP_006553_2_transcription_factor_LBX1__Homo_sapiens_
NP_001072559_1_transcription_factor_LBX1__Xenopus_tropicalis_
NP_001089192_1_transcription_factor_LBX1__Xenopus_laevis_
ACI26672_1_Lbx_homeodomain_protein__Capitella_teleta_
NP_524434_2_ladybird_late__isoform_A__Drosophila_melanogaster_
NP_034822_1_transcription_factor_LBX2__Mus_musculus_

**FigureS1.5:**

**A** bHLH protein phylogenetic trees calculated with ngphylogeny.fr using the FastME/OneClick method, with 100 bootstrap calculation. Note that cnidarians do not have homologs of the MyoD-MRF family, but have significantly expanded independently (orange box). and **B** Nk-family phylogenetic trees calculated with ngphylogeny.fr using the PhyML /OneClick method, with 100 bootstrap calculation. A Cnidarian-specific radiation of VND-related NK2.2 genes is evident (orange box). Nematostella sequences are highlighted in red.

Figure S2.1:

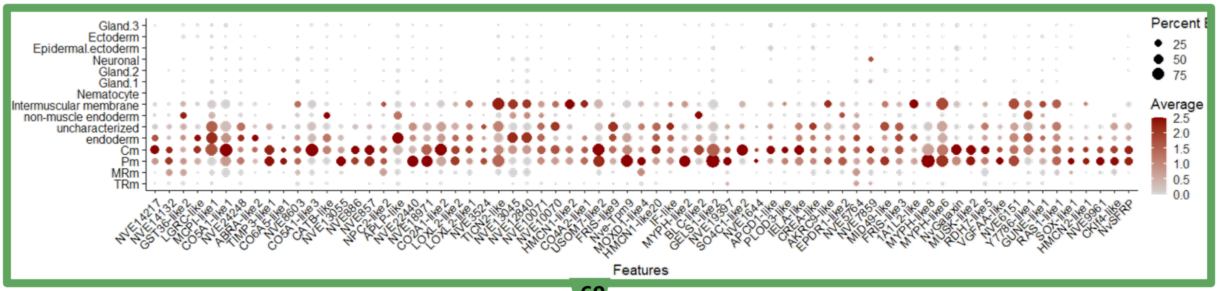


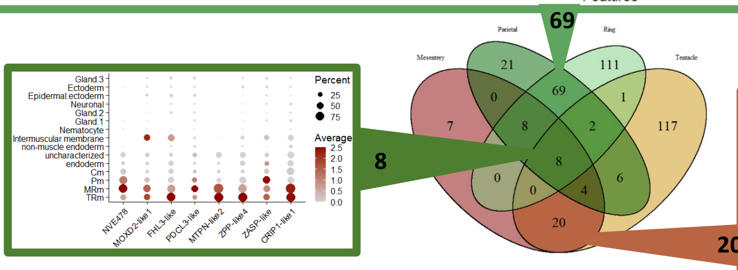


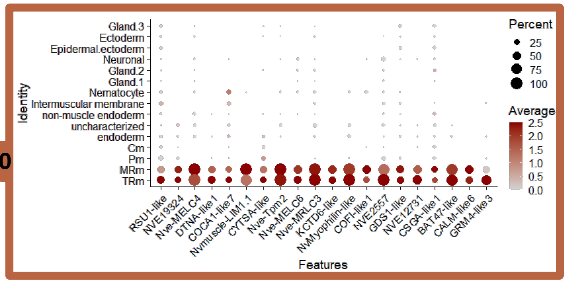


### Supplementary files: Gene lists

Ext.Data1.4

ExtendedData4MUSCLE_StructuralMuscle.csv

**Figure S2.1:**
Set of all genes expressed within any of the differentiated muscle cell populations, but largely absent from the remaining ectodermal cells of the full dataset. Genes were selected as having at least 10 reads associated with any muscle cell cluster from the data subset, but less than 200 reads from the remaining ectodermally-derived dataset. Many genes of the slow muscle group show extensive expression also within the other non-muscle endodermal cell clusters. The central VennDiagram shows the overlapping components, whose expression profiles are illustrated as DotPlots on the full dataset.

Figure S2.2 :

Bodywall set:

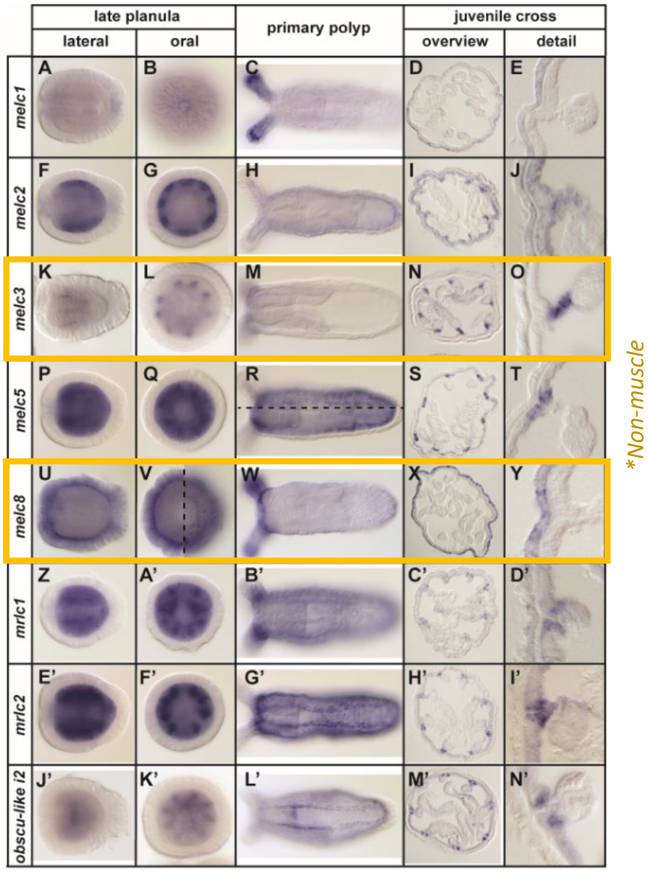


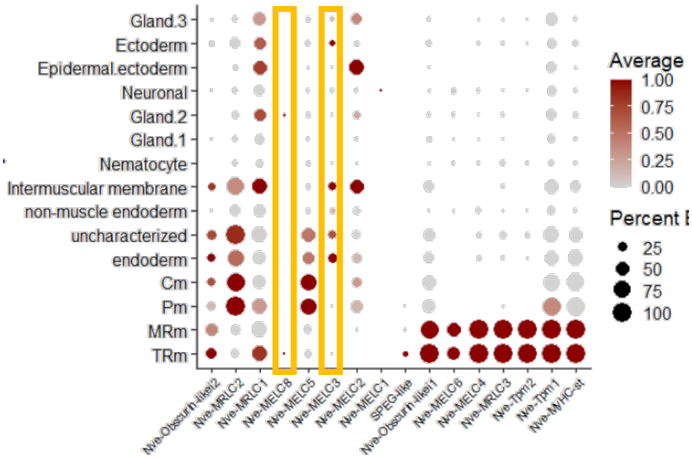


Retractor set:

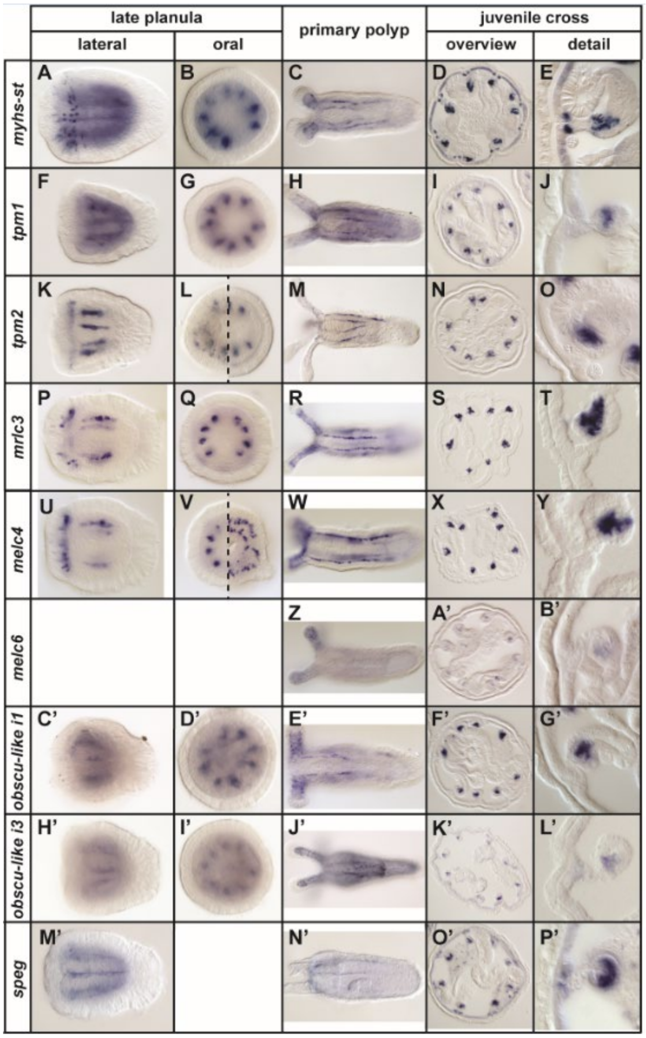

**Figure S2.2:**

Spatial expression profiles for selected structural genes by colorimetric in situ hybridization. Genes representative of the bodywall (slow) gene set are shown on the left. The distribution of each gene within the scRNA dataset is shown in the center. Two genes (yellow bars) are paralogs NOT expressed within differentiated muscle tissue. Genes representative of the fast-contracting retractor muscle gene set are shown in the right panel. In all cases the spatial restriction to the retractor muscles as indicated by the single cell data is confirmed.

Figure S2.3 :

A

**Tentacle Retraction**

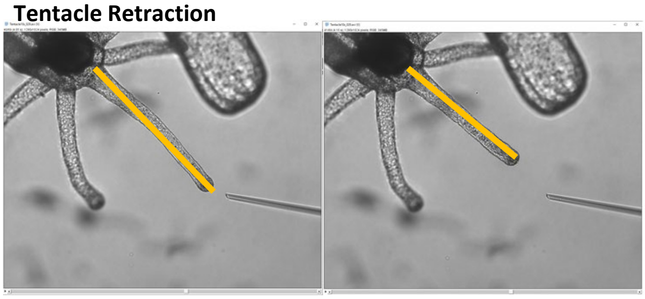

B

**Peristaltic Contraction (CM)**

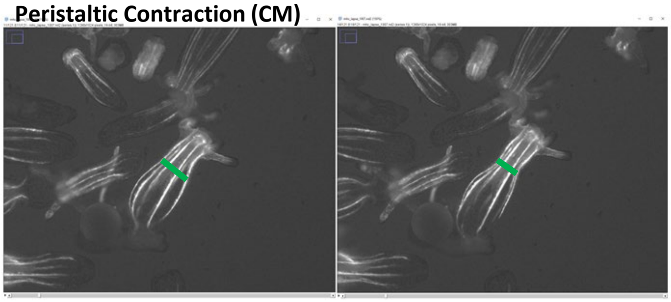

C **Body Retraction (mesentery retractor)**

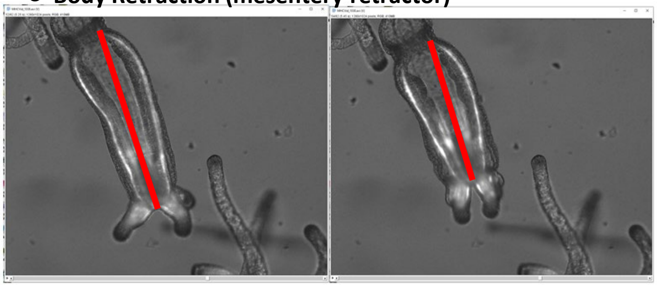

D

measuring the movement of two MHC filaments may yield in a measurement of only one direction of movement since again the animal is 3d but the spatial orientation of the filaments can not be discerned from this view

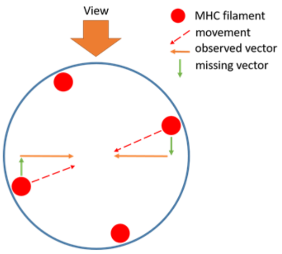


Supplementary files: RawData

ExtendedData5_**SpeedData.xlsx**
ExtendedData6_**VIDEOS**

**Figure S2.3:**
Illustration of image frame measurements for calculation of contraction rates as shown in figure 2D. The measured vector is indicated for all three measurement types: tentacle retraction (ochre), retraction of the body column (red) as a proxy for the mesentery retractor, and body width as a proxy for peristaltic contraction (green). The distance used as circumferential contraction distance is calculated as indicated in **D**

Figure S2.4

A

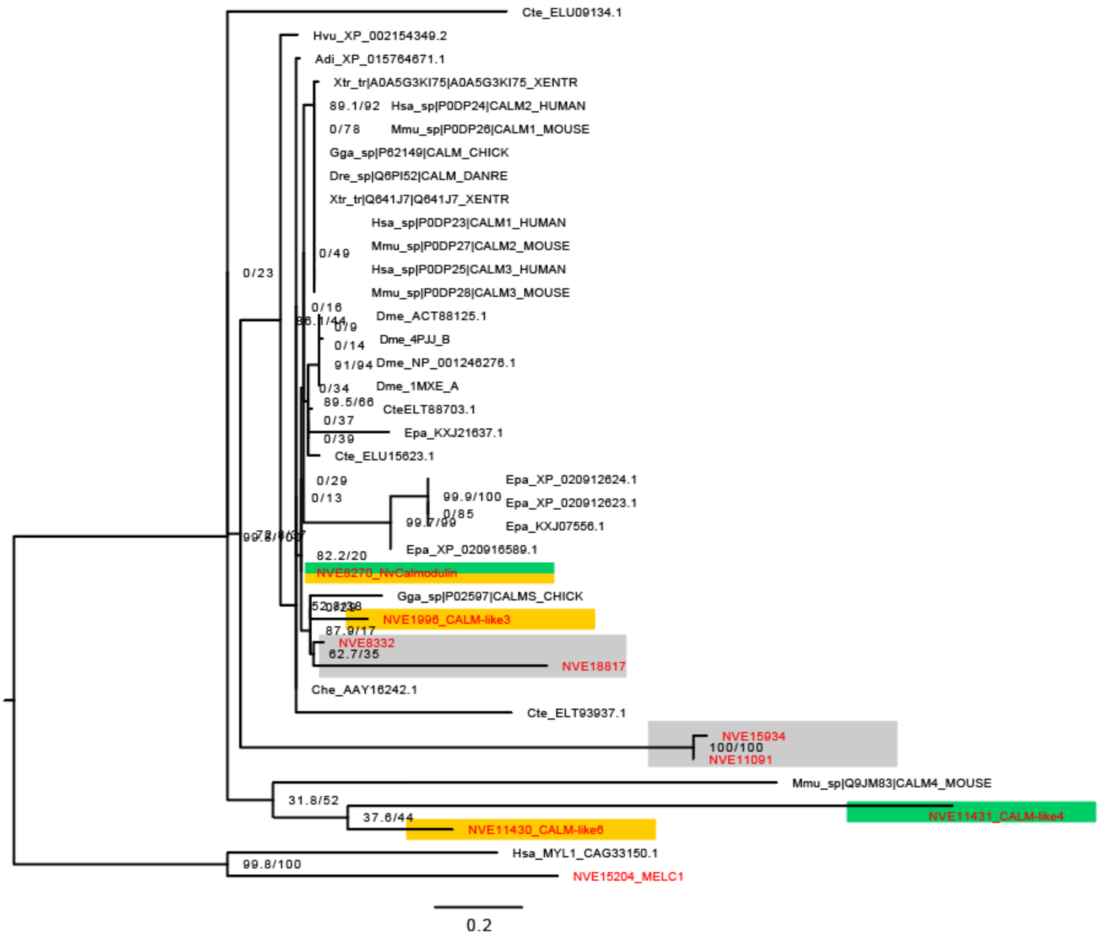


B

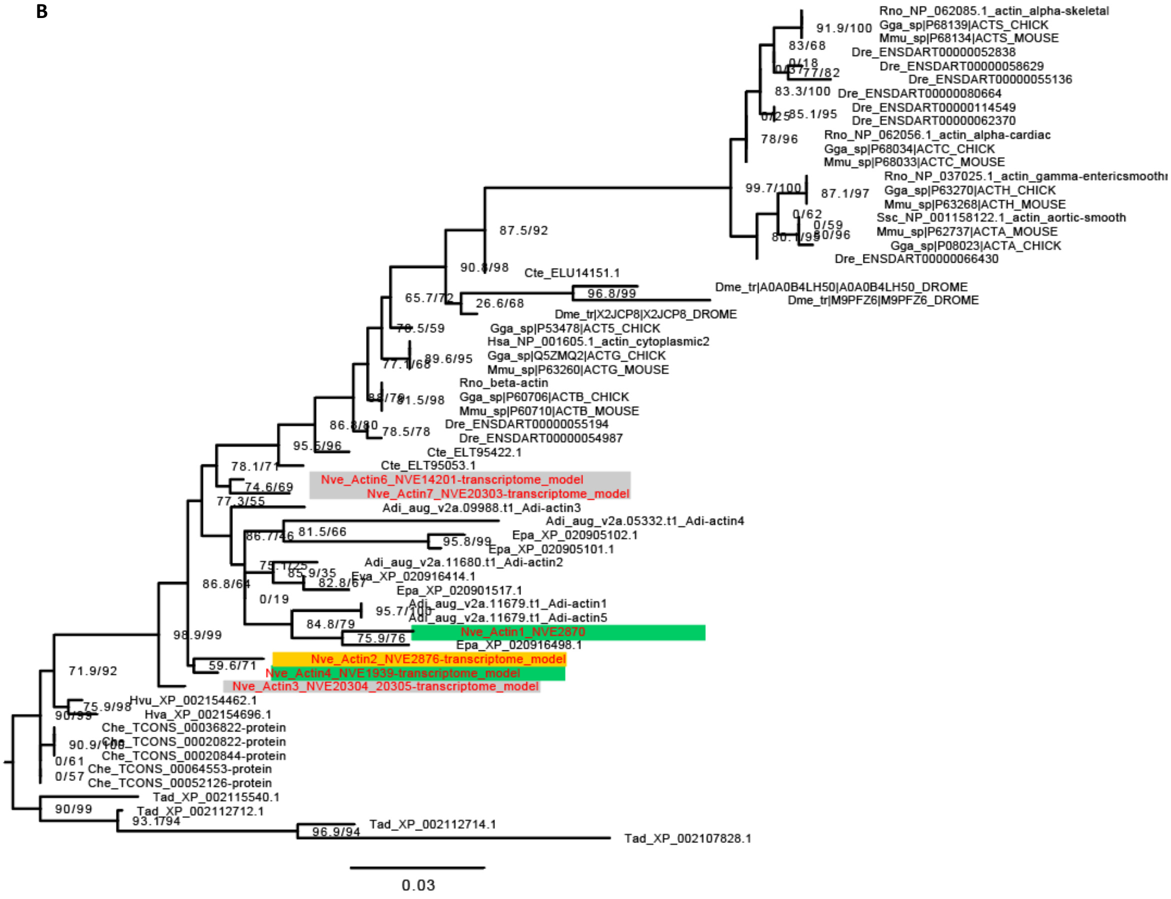

Figure S2.4

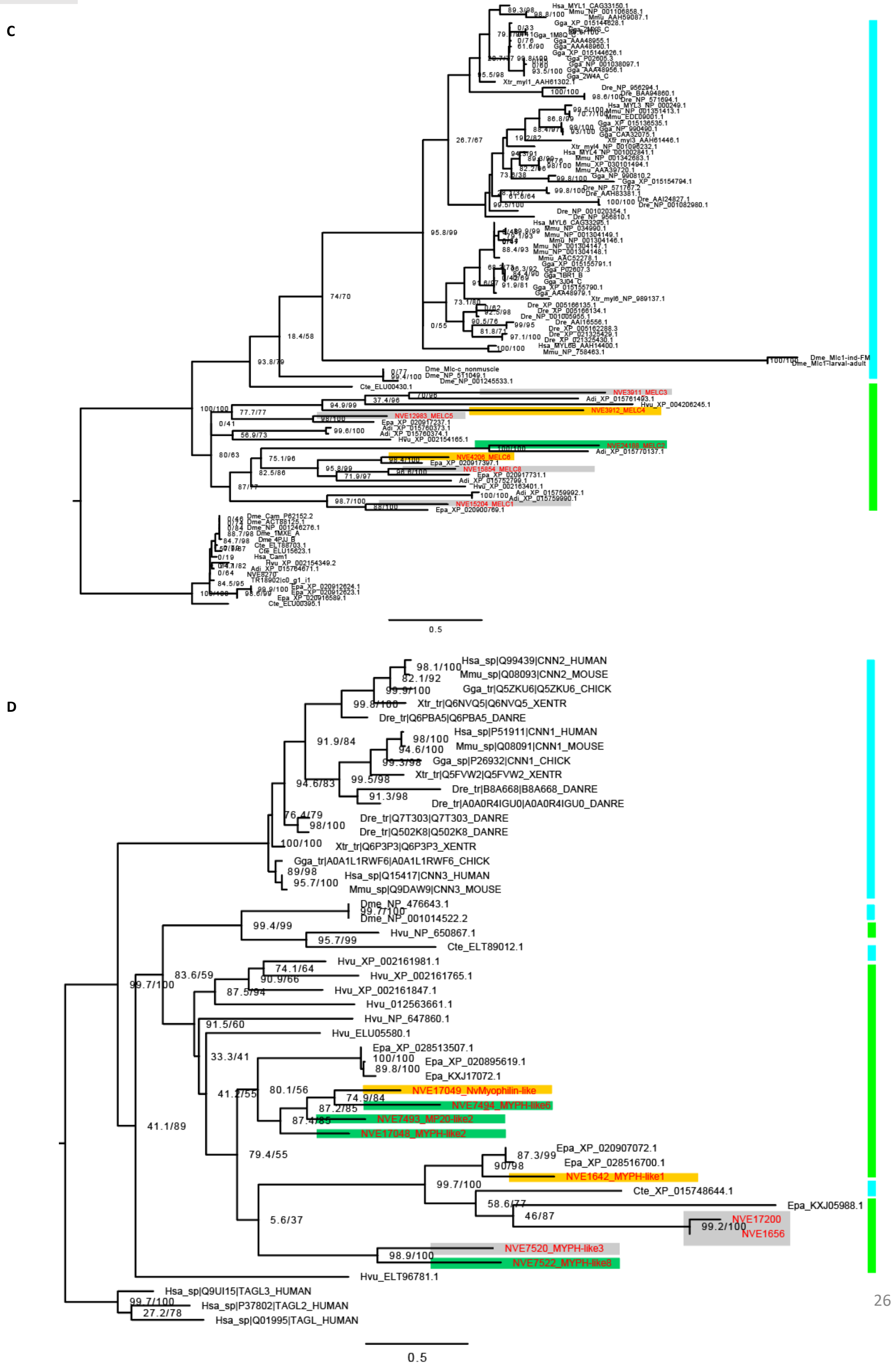
C
D
Hsa_sp|Q99439|CNN2_HUMAN
Mmu_sp|Q08093|CNN2_MOUSE
Gga_tr|Q5ZKU6|Q5ZKU6_CHICK
Xtr_tr|Q6NVQ5|Q6NVQ5_XENTR
Dre_tr|Q6PBA5|Q6PBA5_DANRE
Hsa_sp|P51911|CNN1_HUMAN
Mmu_sp|Q08091|CNN1_MOUSE
Gga_sp|P26932|CNN1_CHICK
Xtr_tr|Q5FVW2|Q5FVW2_XENTR
Dre_tr|B8A668|B8A668_DANRE
Dre_tr|A0A0R4IGU0|A0A0R4IGU0_DANRE
Dre_tr|Q7T303|Q7T303_DANRE
Dre_tr|Q502K8|Q502K8_DANRE
Xtr_tr|Q6P3P3|Q6P3P3_XENTR
Gga_tr|A0A1L1RWF6|A0A1L1RWF6_CHICK
Hsa_sp|Q15417|CNN3_HUMAN
Mmu_sp|Q9DAW9|CNN3_MOUSE
Dme_NP_476643.1
Dme_NP_001014522.2
Hvu_NP_650867.1
Cte_ELT89012.1
Hvu_XP_002161981.1
Hvu_XP_002161765.1
Hvu_XP_002161847.1
Hvu_012563661.1
Hvu_NP_647860.1
Hvu_ELU05580.1
Epa_XP_028513507.1
Epa_XP_020895619.1
Epa_KXJ17072.1
NVE17049_NvMyophilin-like
NVE7494_MYPH-like6
NVE7493_MP20-like2
NVE17048_MYPH-like2
Epa_XP_020907072.1
Epa_XP_028516700.1
NVE1642_MYPH-like1
Cte_XP_015748644.1
Epa_KXJ05988.1
NVE17200
NVE1656
NVE7520_MYPH-like3
NVE7522_MYPH-like8
Hvu_ELT96781.1
Hsa_sp|Q9UI15|TAGL3_HUMAN
Hsa_sp|P37802|TAGL2_HUMAN
Hsa_sp|Q01995|TAGL_HUMAN
0.5

Figure S2.4

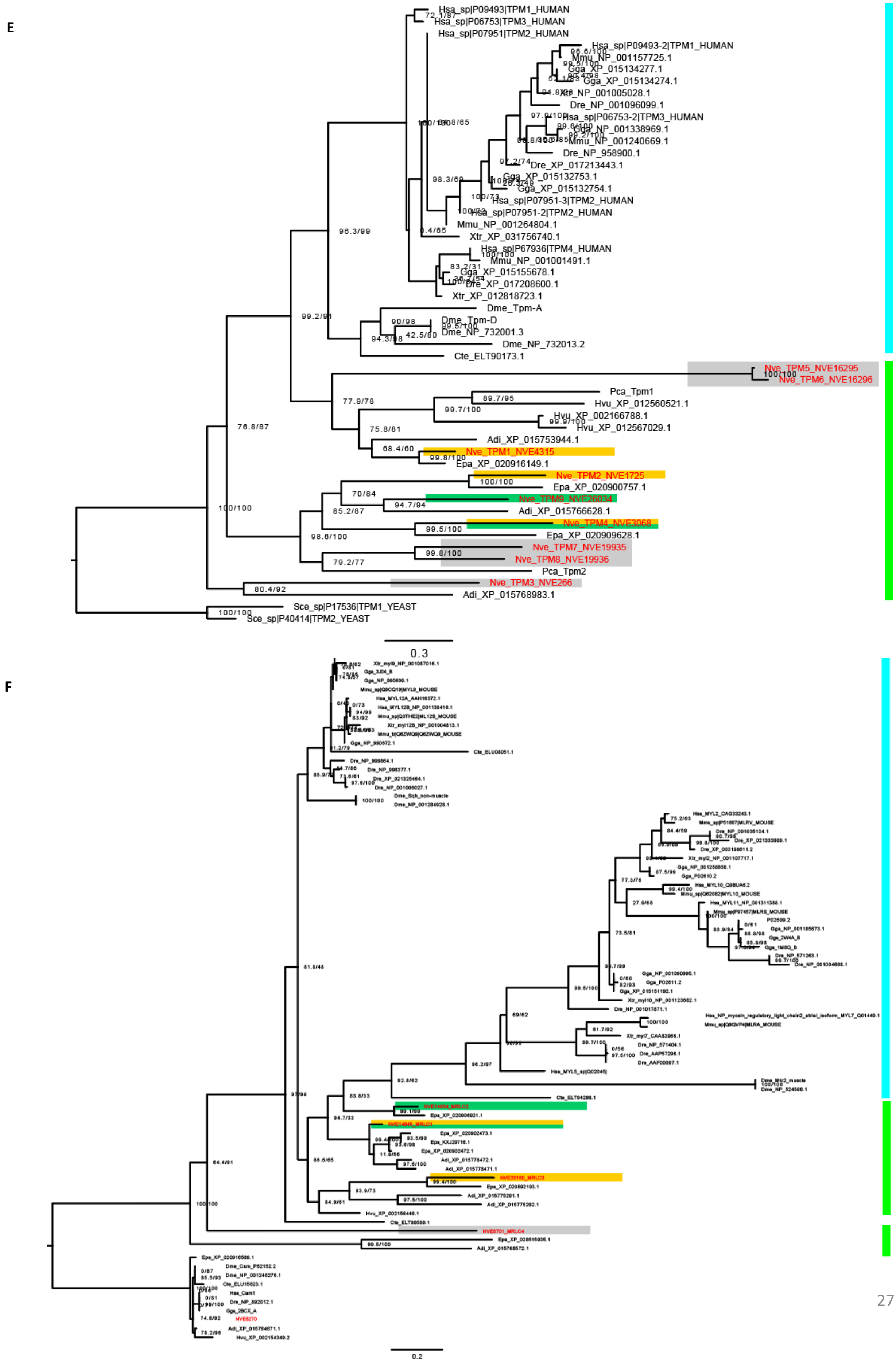
E
Hsa_sp|P09493|TPM1_HUMAN
Hsa_sp|P06753|TPM3_HUMAN
Hsa_sp|P07951|TPM2_HUMAN
Hsa_sp|P09493-2|TPM1_HUMAN
Mmu_NP_001157725.1
Gga_XP_015134277.1
Gga_XP_015134274.1
Xtr_NP_001005028.1
Dre_NP_001096099.1
Hsa_sp|P06753-2|TPM3_HUMAN
Gga_NP_001338969.1
Mmu_NP_001240669.1
Dre_NP_958900.1
Dre_XP_017213443.1
Gga_XP_015132753.1
Gga_XP_015132754.1
Hsa_sp|P07951-3|TPM2_HUMAN
Hsa_sp|P07951-2|TPM2_HUMAN
Mmu_NP_001264804.1
Xtr_XP_031756740.1
Hsa_sp|P67936|TPM4_HUMAN
Mmu_NP_001001491.1
Gga_XP_015155678.1
Dre_XP_017208600.1
Xtr_XP_012818723.1
Dme_Tpm-A
Dme_Tpm-D
Dme_NP_732001.3
Dme_NP_732013.2
Cte_ELT90173.1
Nve_TPM5_NVE16295
Nve_TPM6_NVE16296
Pca_Tpm1
Hvu_XP_012560521.1
Hvu_XP_002166788.1
Hvu_XP_012567029.1
Adi_XP_015753944.1
Nve_TPM1_NVE4315
Epa_XP_020916149.1
Nve_TPM2_NVE1725
Epa_XP_020900757.1
Nve_TPM9_NVE26034
Adi_XP_015766628.1
Nve_TPM4_NVE3068
Epa_XP_020909628.1
Nve_TPM7_NVE19935
Nve_TPM8_NVE19936
Pca_Tpm2
Nve_TPM3_NVE266
Adi_XP_015768983.1
Sce_sp|P17536|TPM1_YEAST
Sce_sp|P40414|TPM2_YEAST
96.3/99
99.2/91
77.9/78
89.7/95
99.7/100
75.8/81
68.4/60
76.8/87
100/100
70/84
94.7/94
85.2/87
99.5/100
98.6/100
99.8/100
79.2/77
80.4/92
0.3
F
0.2

**Figure S2.4:**

Gene trees for paralogous structural genes Expression profiles in terms of muscle type are indicated as boxes (Fast muscle: orange; Slow muscle: green; non-muscle: grey). All *Nematostella* gene models are highlighted in red. Colored boxes along the right side indicate the distribution of cnidarian (bright green) vs. bilaterian (cyan) sequences. Nodes are labeled with SH-aLRT /ultrafast bootstrap support (%). **A** Calmodulin **B** Actin **C** MELC **D** Calponin **E** MRLC **F** Tropomyosin. *Aqu: Amphimedon queenslandica, Adi: Acropora digitifera, Hvu: Hydra vulgaris, Epa: Exaiptasia pallida, Cte: Capitella teleta, Dme: Drosophila melanogaster, Che: Clytia hemisphaerica, Mmu: Mus musculus, ga: Gallus gallus*, *Xtr: Xenopus tropicalis, Hsa: Homo sapiens, Dre: Danio rerio*

Figure S3

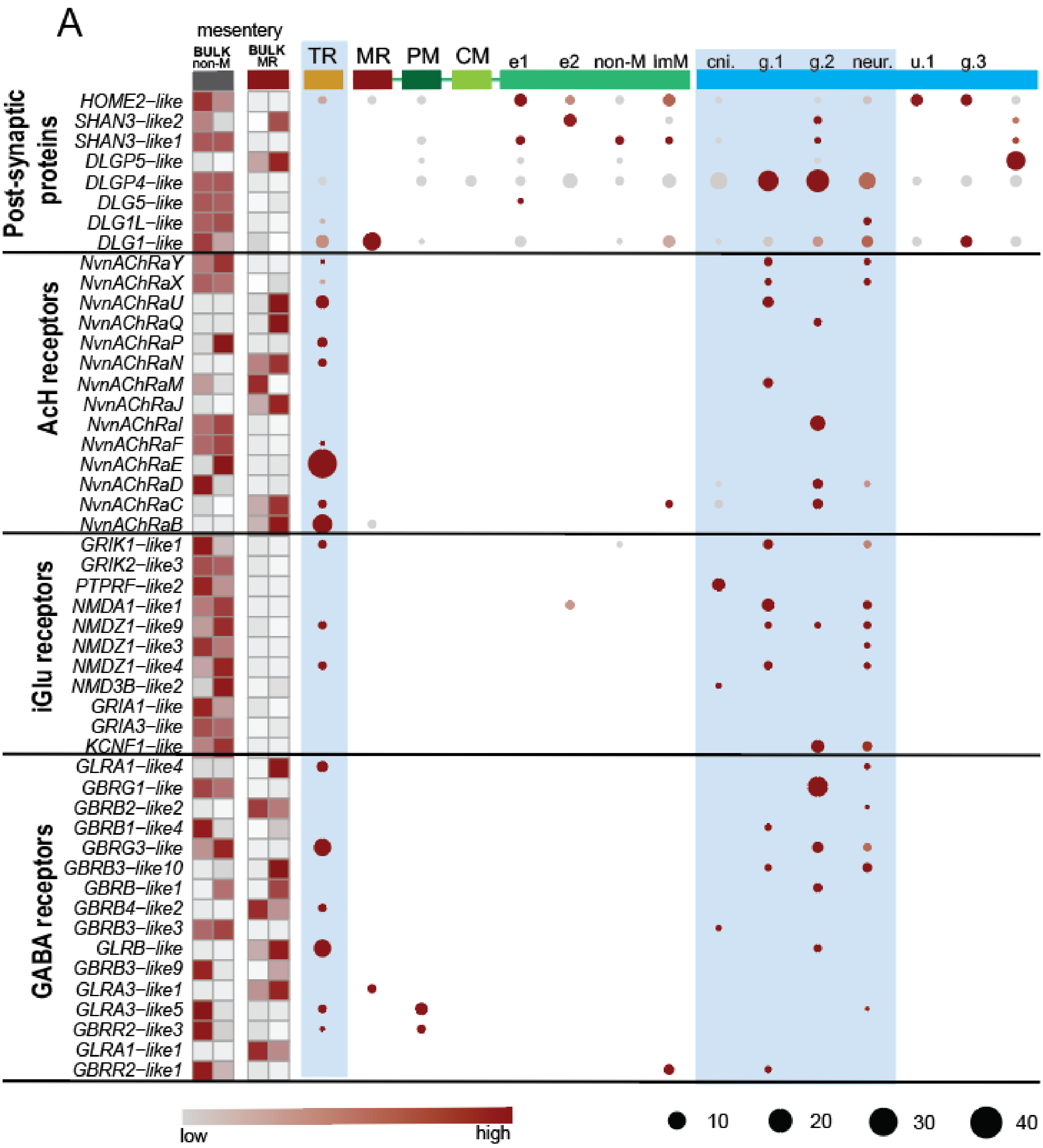
A
mesentery
BULK non-M
BULK MR
TR
MR
PM
CM
e1
e2
non-M
ImM
cni.
g.1
g.2
neur.
u.1
g.3
Post-synaptic proteins
HOME2–like
SHAN3–like2
SHAN3–like1
DLGP5–like
DLGP4–like
DLG5–like
DLG1L–like
DLG1–like
AcH receptors
NvnAChRaY
NvnAChRaX
NvnAChRaU
NvnAChRaQ
NvnAChRaP
NvnAChRaN
NvnAChRaM
NvnAChRaJ
NvnAChRaI
NvnAChRaF
NvnAChRaE
NvnAChRaD
NvnAChRaC
NvnAChRaB
iGlu receptors
GRIK1–like1
GRIK2–like3
PTPRF–like2
NMDA1–like1
NMDZ1–like9
NMDZ1–like3
NMDZ1–like4
NMD3B–like2
GRIA1–like
GRIA3–like
KCNF1–like
GABA receptors
GLRA1–like4
GBRG1–like
GBRB2–like2
GBRB1–like4
GBRG3–like
GBRB3–like10
GBRB–like1
GBRB4–like2
GBRB3–like3
GLRB–like
GBRB3–like9
GLRA3–like1
GLRA3–like5
GBRR2–like3
GLRA1–like1
GBRR2–like1
low
high
10
20
30
40

**FigureS3:**
Relative expression plot of putative post-synaptic proteins and ionotrophic receptors, organized as in Figure 2A. Most ionic receptor proteins are found within ectodermal derivatives (blue highlight). Acetylcholine subuntis are detected within the BULK dataset, but are largely undetectable from the endodermal cell populations.

**Movie S1.**

Example time-lapse film of MHC::mCherry line used to calculate contraction speeds as shown in Figure 2D. Measurements from single frames are found in Extended Data S2

**Extended Data S1 (separate file)**

Spreadsheets of gene lists derived from different analysis approaches. **ExtDataS1.1**: list of upregulated genes in the BULK dataset shown in Figure 1D. **ExtDataS1.2**: output of FindAllMarkers function of the Seurat Vs3 R package, for the clusters identified in Figure 1C. **ExtDataS1.3**: output of FindAllMarkers function of the Seurat Vs3 R package, for the clusters identified in Figure 1E, excluding the set of all putative Transcription Factors. Expression profiles are imaged in Figure S1.3E. **ExtDataS1.4**: output of FindAllMarkers function of the Seurat Vs3 R package, for the clusters identified in Figure 1E from the set of all putative Transcription Factors. Expression profiles are imaged in Figure S1.4A **ExtDataS1.5**: list of all putative transcription factors detected in any differentiated muscle cell population, but absent from the ectodermal derivatives of the full dataset, as imaged in Figure S1.4B. The intersection between the lists generated for each muscle cell type is illustrated in Figure S1.4C. **ExtDataS1.6**: list of genes detected in any differentiated muscle cell population, excluding the set of all putative transcription factors, and absent from the ectodermal derivatives of the full dataset, as imaged in Figure S2.1.

**Extended Data S2 (separate file)**

Spreadsheet of measurements used to calculate the muscle contraction speed shown in Figure 2D.

**Extended Data S3 (separate file)**

Cross reference between gene models and annotations.

**Extended Data S4 (separate file)**

R script for generating the analysis presented in this manuscript.

**Extended Data S5 (separate file)**

Protein alignments used to generate trees in Figures S1.5 and S2.4.
